# Activity-Dependent Modulation of Synapse-Regulating Genes in Astrocytes

**DOI:** 10.1101/2020.12.30.424365

**Authors:** I Farhy-Tselnicker, MM Boisvert, H Liu, C Dowling, GA Erikson, E Blanco-Suarez, C Farhy, M Shokhirev, JR Ecker, NJ Allen

**Affiliations:** Molecular Neurobiology Laboratory, Salk Institute for Biological Studies, 10010 N Torrey Pines Rd, La Jolla, CA, 92037, USA; Genomic Analysis Laboratory, Salk Institute for Biological Studies, 10010 N Torrey Pines Rd, La Jolla, CA, 92037, USA; Razavi Newman Integrative Genomics and Bioinformatics Core, Salk Institute for Biological Studies, 10010 N Torrey Pines Rd, La Jolla, CA, 92037, USA; Howard Hughes Medical Institute, Salk Institute for Biological Studies, 10010 N Torrey Pines Rd, La Jolla, CA, 92037, USA; Sanford Burnham Prebys Medical Discovery Institute, 10901 N Torrey Pines Rd, La Jolla, CA, 92037, USA; Department of Biology, Texas A&M University, 301 Old Main drive, College Station, TX, 77843, USA; Jungers Center for Neuroscience Research, Department of Neurology, Oregon Health and Science University, Portland, OR, 97239, USA; Department of Neurosurgery, Thomas Jefferson University Hospital for Neuroscience, 900 Walnut St, Philadelphia, PA, 19107, USA

**Keywords:** Astrocytes, synapse development, gene expression, neuronal activity, visual cortex

## Abstract

Astrocytes regulate the formation and function of neuronal synapses via multiple signals, however, what controls regional and temporal expression of these signals during development is unknown. We determined the expression profile of astrocyte synapse-regulating genes in the developing mouse visual cortex, identifying astrocyte signals that show differential temporal and layer-enriched expression. These patterns are not intrinsic to astrocytes, but regulated by visually-evoked neuronal activity, as they are absent in mice lacking glutamate release from thalamocortical terminals. Consequently, synapses remain immature. Expression of synapse-regulating genes and synaptic development are also altered when astrocyte signaling is blunted by diminishing calcium release from astrocyte stores. Single nucleus RNA sequencing identified groups of astrocytic genes regulated by neuronal and astrocyte activity, and a cassette of genes that show layer-specific enrichment. Thus, the development of cortical circuits requires coordinated signaling between astrocytes and neurons, identifying astrocytes as a target to manipulate in neurodevelopmental disorders.

## Introduction

Synapses are points of contact where electro-chemical signals are transferred between neurons in a given circuit (Petzoldt and Sigrist, 2014; Südhof, 2018). In many brain regions, such as the mammalian visual cortex (VC), the majority of synapses are contacted by a process from an astrocyte, a major type of glial cell (Bernardinelli et al., 2014a; Genoud et al., 2006; Ventura and Harris, 1999). Synapse development is a complex multi-step process (Allen, 2013; Batool et al., 2019). In the VC, synapses begin to form at around postnatal day (P) 7, peak at P14, and stabilize towards P28, remaining stable to adulthood (Blue and Parnavelas, 1983a, b; Farhy-Tselnicker and Allen, 2018; Li et al., 2010). Within this time period numerous developmental programs are being executed from the molecular to the behavioral levels. Astrocytes appear in the cortex at birth, and populate the cortex, migrating, proliferating and maturing throughout the first postnatal month, coincidently with synapse development (Farhy-Tselnicker and Allen, 2018). Synaptic deficits, for example caused by mutations in synapse-related genes expressed in either neurons or astrocytes, are associated with developmental disorders such as autism spectrum disorder and epilepsy (Lepeta et al., 2016). Therefore, understanding how synaptic development is regulated will provide important insights into how circuits form in health and misfunction in disease.

In the past 20 years many astrocyte-secreted factors have been identified that regulate distinct stages of excitatory glutamatergic synapse formation and maturation (Baldwin and Eroglu, 2017). For example, thrombospondin family members induce formation of structurally normal but functionally silent synapses (Christopherson et al., 2005; Eroglu et al., 2009). Glypicans induce the formation of active synapses by recruiting GluA1 to the postsynaptic side (Allen et al., 2012; Farhy-Tselnicker et al., 2017) and chordin-like 1 induces synapse maturation by recruiting GluA2 to the postsynaptic side (Blanco-Suarez et al., 2018). The majority of these signals have been identified using *in vitro* cell culture approaches and analyzed at distinct ages and across brain regions *in vivo*. To understand how these diverse signals act together to regulate formation of a complete circuit, it is important to determine when and where each of them is expressed *in vivo*, and how its expression correlates with the distinct stages of synaptic development that it regulates. Furthermore, the regulatory mechanisms that control the developmental expression level of these astrocyte synapse-regulating genes are largely unknown.

We chose to address these questions by analyzing the *in vivo* development of both astrocytes and synapses in the mouse visual cortex (VC). The rodent VC is composed of heterogeneous populations of neurons, approximately 80% excitatory glutamatergic and 20% inhibitory GABAergic (Markram et al., 2004). Glutamatergic neurons are arranged in spatially defined layers, with distinct connectivity patterns between layers, as well as with other cortical and subcortical regions (Bannister, 2005; Douglas and Martin, 2004). The main subcortical region that projects to the VC is the lateral geniculate nucleus of the thalamus (LGN), relaying visual information received from the retina. Although thalamic projections arrive at the cortex before birth, synapses develop postnatally (Li et al., 2010; Lopez-Bendito and Molnar, 2003). Before eye-opening (from birth to ~ P12 in mice), spontaneous retinal activity evokes correlated cortical responses (Gribizis et al., 2019; Hanganu et al., 2006) that are important for the correct establishment of thalamo-cortical synapses (Cang et al., 2005). Eye-opening marks a step towards synapse maturation in the VC, with the appearance of visually-evoked neuronal responses across the retinal-LGN-VC circuit (Espinosa and Stryker, 2012; Hooks and Chen, 2006, 2020). Preventing this step from taking place by methods of visual deprivation, such as rearing animals in the dark, has been shown to delay synaptic maturation in the VC of several species including mice at the transcriptomic (Hsu et al., 2018; Majdan and Shatz, 2006; Tropea et al., 2006), structural (Albanese et al., 1983; Freire, 1978) and functional levels (Desai et al., 2002; Funahashi et al., 2013; Ishikawa et al., 2014; Ko et al., 2014), as well as perturb structural astrocyte maturation (Müller, 1990; Stogsdill et al., 2017). This well-characterized process of experience dependent synaptic development and maturation makes the VC an ideal place to investigate the role of astrocytes in regulating the different stages of synaptogenesis. Recent work has shown that similarly to neurons, cortical astrocytes are also spatially arranged in diverse populations (Batiuk et al., 2020; Bayraktar et al., 2020; John Lin et al., 2017; Lanjakornsiripan et al., 2018), in line with evidence from other brain regions showing astrocyte heterogeneity (Chaboub and Deneen, 2012; Chai et al., 2017; Khakh and Deneen, 2019; Oberheim et al., 2012; Rusnakova et al., 2013; Schitine et al., 2015). However, whether this astrocyte diversity has any impact on their regulation of synapse formation or maturation in the VC, is unknown.

Here we use RNA sequencing and *in situ* hybridization to obtain the developmental transcriptome of astrocytes in the mouse VC *in vivo*. We find that astrocyte synapse-regulating genes display differential temporal and spatial expression patterns which correspond to stages of synapse initiation and maturation. Furthermore, we find that developmental regulation of these genes, namely glypican 4 and chordin-like 1, depends on visually evoked neuronal activity, with additional regulation by astrocyte Ca^2+^ activity. Manipulating either neuronal or astrocytic activity leads to shifts in synaptic development and maturation. Finally, single-nucleus RNA sequencing analysis reveals diverse populations of astrocytes in the developing VC, as well as identifies novel groups of genes that are regulated by neuronal and astrocyte activity. These findings demonstrate how astrocyte expression of synapse-regulating genes is controlled during development, and how synapse maturation is dependent on neuron-astrocyte communication. These data further provide an important resource for future studies of astrocyte development and astrocyte regulation of synapse formation.

## Results

### Development of astrocytes and synapses in the mouse visual cortex

To determine how synapse-regulating genes in astrocytes participate in synaptic development we first analyzed the development of astrocytes and synapses in the mouse visual cortex (VC) over the first postnatal month, focusing on ages correlating with stages of astrocyte and synapse development: postnatal day (P) 1, astrocytes are being born; P4, astrocytes continue to expand in the cortex, synapses not present; P7, synapse initiation; P14, start of synapse maturation; P28, stable synapses (Fig 1A,C). To provide spatial as well as temporal information to this data, all analyses were conducted separately in each cortical layer (L1, L2/3, L4, L5, L6) (Fig 1B,C).

**Figure 1.**
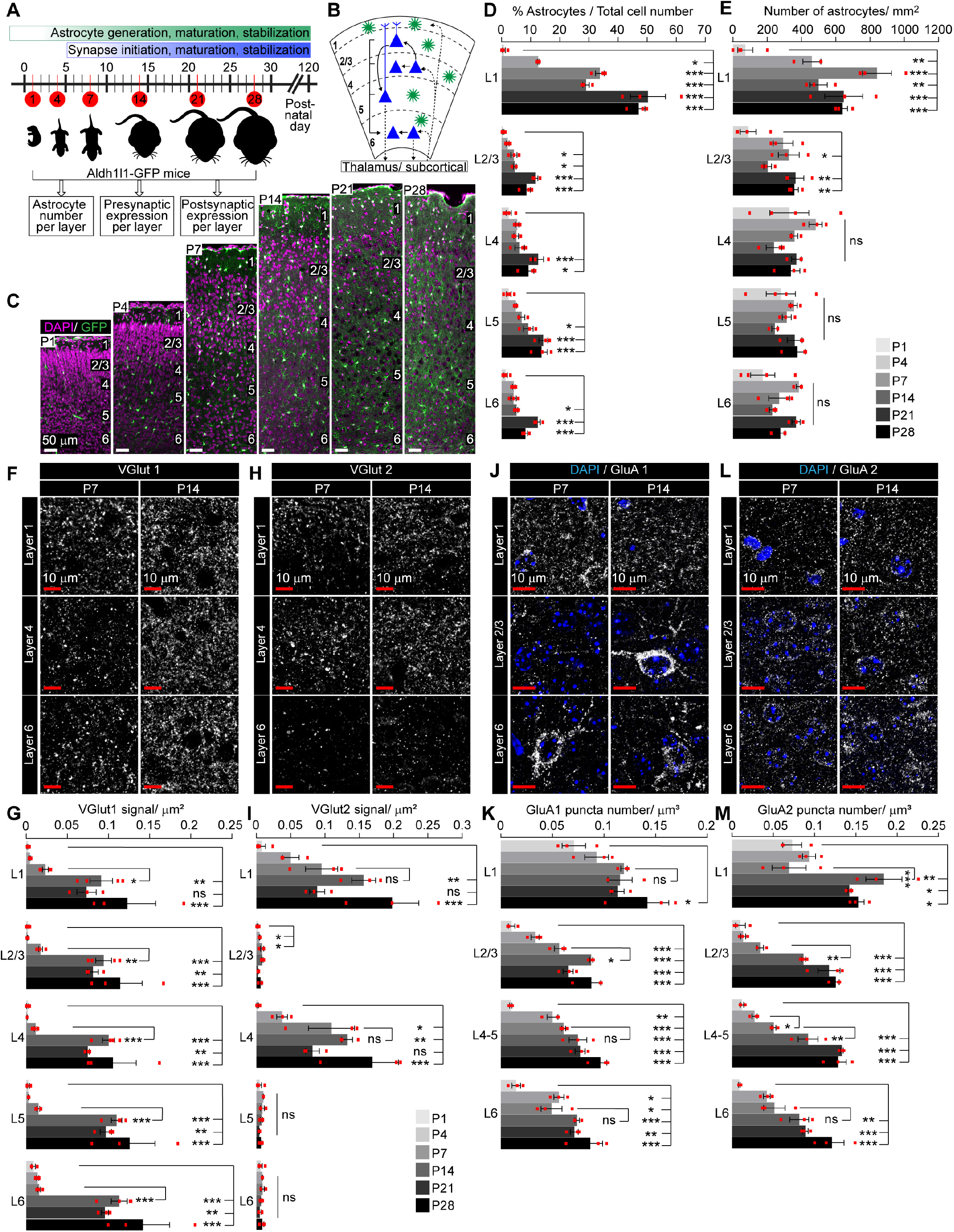
Development of astrocytes and synapses in the mouse visual cortex. See also Fig S1, Table S1. **A.** Schematic of time points of VC development analyzed, corresponding to synapse development. **B.** Diagram of VC depicting neuronal (blue) laminar arrangement and connectivity (arrows). Astrocytes (green) are present in all layers. **C-E.** Astrocyte number increases in the VC across development. **C**. Example images of the VC from Aldh1l1-GFP mice at time points analyzed. GFP marks astrocytes (green), DAPI (magenta) labels nuclei. Layers labeled by numbers on the right in each panel. **D.** Quantification of C, astrocytes as a percentage of total cells within each cortical layer. **E.** Quantification of C, number of astrocytes per mm^2^ of VC within each layer. **F-I.** Developmental expression pattern of the presynaptic proteins VGlut1 and VGlut2 in each cortical layer. **F, H.** Example images of VGlut1 or VGlut2 protein levels (white puncta). **G, I.** Quantification of F, H, density of VGlut1 or VGlut2 signal as threshold area per μm^2^. **J-M.** Developmental expression pattern of the postsynaptic AMPAR subunits GluA1 and GluA2 within each cortical layer. **J, L.** Example images of GluA1 or GluA2 protein levels (white puncta), DAPI (blue) labels nuclei. **K, M.** Quantification of J, L number of GluA1 or GluA2 positive puncta per cortical volume (μm^3^). Scale bars in C: 50μm; In F, H, J, L: 10μm. In C-E: N=4 mice for P1; N=3 mice for P4-P28. In F-M: N= 3 mice/ age. Graphs show mean ±s.e.m., red squares average of individual mouse. *P ≤ 0.05 **P<0.01, ***P<0.001, ns (not significant) by one-way ANOVA comparing expression between time points within each layer.

Since astrocytes are still migrating and dividing during this time we first sought to quantify how astrocyte density and/or the fraction of astrocytes relative to all other cells change with development. To do this we utilized the Aldh1l1-GFP mouse line (Dougherty et al., 2010; Tien et al., 2012), where astrocytes express GFP under the Aldh1l1 promoter. This line has been previously used to study astrocytes across development (John Lin et al., 2017; Stogsdill et al., 2017). Immunostaining of brain sections from Aldh1l1-GFP mice at P7 and P28 with antibodies against known astrocyte markers Aldh1l1, S100β, and Sox9 showed high overlap between GFP and marker positive cells (Fig S1D-F), further validating its usage. Close to birth (P1) very few astrocytes are present, comprising 0.5-2% of the total cell number in the VC (GFP positive cells as a percentage of all cells marked by the nuclear dye DAPI), with a significantly higher percentage of astrocytes in deeper layers than upper layers (Fig 1C-D, Table S1). The astrocyte percentage increases with development in all cortical layers, peaking at P21-P28. At this time astrocytes are ~10% of total cell number in L2-6, and ~ 50% in L1 (Fig 1C,D, Table S1). As the brain grows during the first postnatal month, the distance between cells grows to accommodate the increase in cell size and complexity, as evident by a significant decrease in DAPI positive nuclei per mm^2^ that occurs from P1 to P14-P28 (Fig S1A, Table S1). Despite this decrease in total cell density, the density of astrocytes remains constant in all cortical layers and across ages, with the exception of L1-2/3, where astrocyte density is significantly lower at P1 (Fig 1C,E). This stability in astrocyte density is likely explained by new astrocytes still being generated in the weeks after birth (Ge et al., 2012).

We next asked how synaptic proteins change across development within cortical layers to correlate with astrocyte development. We focused on glutamatergic synapses, as the majority of thus far identified astrocyte-expressed synapse-regulating factors are known to affect these synapses (Allen, 2013; Baldwin and Eroglu, 2017; Bosworth and Allen, 2017). To detect presynaptic terminals we stained with VGlut1 to identify local cortico-cortical connections (Fig 1F,G, Fig S1B, Table S1), and VGlut2 to identify thalamo-cortical connections (Fig 1H,I, Fig S1C, Table S1) (Fremeau et al., 2001). To detect postsynaptic AMPA glutamate receptors we stained for GluA1 subunits typically associated with immature synapses, and GluA2 subunits associated with mature synapses (Brill and Huguenard, 2008; Kumar et al., 2002) (Fig 1J-M, Table S1). VGlut1 immunoreactivity greatly increases between P7 and P14 in all cortical layers and remains stable at later ages (Fig 1G, Table S1). VGlut2 levels steadily increase from P1 to P14, and then remain stable. VGlut2 immunoreactivity is significantly higher in L1 and L4 than other layers at all ages, consistent with thalamic innervation zones (Fig 1I, Table S1)(Lopez-Bendito and Molnar, 2003). GluA1 levels increase from P1 to P7 and then remain mostly constant through P28 (Fig 1K, Table S1). GluA2 immunoreactivity significantly increases from P7 to P14 in L1-5, and then remains stable to P28 (Fig 1M, Table S1). At all ages, the levels of GluA1 and GluA2 are significantly higher in L1 than all other layers, with most prominent and statistically significant differences occurring at early time points (at P1-7 AMPAR subunit signal is ~2-7-fold higher in L1 compared to L2-6; at P14-P28 AMPAR signal is 1.2-2-fold higher in L1 compared to L2-6; Table S1).

Taken together this analysis shows that astrocyte numbers increase with postnatal development to represent ~10% of all cells in the VC by P21. Synaptic proteins increase in level across the same time period, with the majority of significant changes occurring between P7 and P14. Spatial analysis revealed astrocyte/synapse-specific patterns. While astrocyte numbers and AMPAR levels show higher abundance in L1, VGlut1 is evenly distributed across all layers. VGlut2 on the other hand is most abundant in L1 and 4 throughout development. Thus, astrocyte and synapse development during the first postnatal month is non-uniform and has specific spatial and temporal programs.

### Determination of the astrocyte transcriptome across visual cortex development

In addition to increasing in number across postnatal development, astrocytes increase in size and morphological complexity (Bushong et al., 2004; Stogsdill et al., 2017) and undergo gene expression changes as shown by microarray studies (Cahoy et al., 2008). To specifically identify how the gene expression of VC astrocytes changes with development we performed RNA sequencing of bulk astrocyte mRNA at postnatal ages correlating to distinct stages of synaptogenesis (P7, P14, P28), as well as adult (P120; Fig 2A), using the previously characterized Ribo-tag method (Rpl22-HA; Gfap-cre73.12 – astrocyte-ribotag)(Boisvert et al., 2018). Similar to our previous analysis at P120, at P28 we found a high overlap between the HA ribosome tag and the astrocyte marker S100β by immunostaining, with minimal overlap with other cell type markers (Fig S2A-C), as well as significant enrichment in astrocyte-specific genes over other cell types in the mRNA isolated by HA immunopurification (IP) compared to total VC mRNA (input) (Fig S2D).

**Figure 2.**
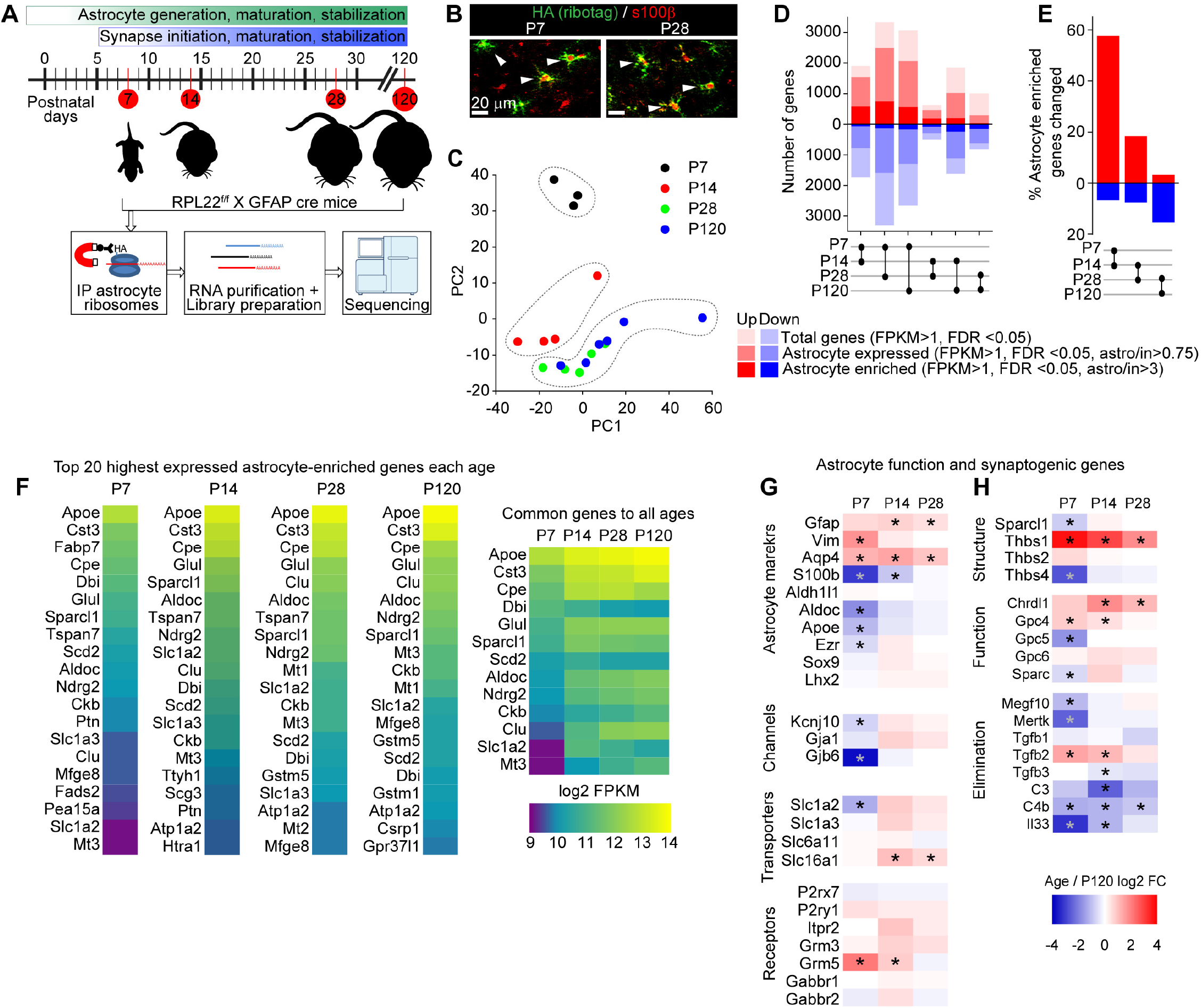
Determination of the astrocyte transcriptome across visual cortex development. See also Fig S2, Table S2. **A.** VCs from Rpl22-HA^f/+^; Gfap-cre 73.12 mice were collected at different time points corresponding to synapse development and maturation and subjected to Ribo-tag pulldown protocol, followed by RNA purification, library preparation and sequencing. **B.** Example images of VC at P7 and P28, showing colocalization between Ribo-tag (green, HA) and astrocyte marker s100β (red). **C.** PC analysis of RNAseq data shows P7 and P14 samples clustering separately from other ages, while P28 and P120 samples cluster together, suggesting similar gene expression profiles (N=3 at P7, 4 at P14, 5 at P28, 3 at P120. For statistical comparisons 3xP120 samples published in (Boisvert et al., 2018) were added to increase the power of the analysis, giving an N=6 P120). Scale bars = 20 μm. **D.** Pairwise comparison of differentially expressed genes (DEGs; red – upregulated, blue - downregulated) between each time point showing total genes (all DEGs identified with FPKM>1), astrocyte expressed genes (expression level in pulldown sample/input >0.75), and astrocyte enriched genes (expression level in pulldown sample/input >3). **E.** Percent of all astrocyte enriched genes that are differentially expressed between each age. **F.** Heatmaps of top 20 astrocyte enriched genes at each age, sorted by expression level, along with 13 genes common to all time points. Colors represent log2 FPKM of expression level. **G, H.** Heatmaps of select genes related to astrocyte function (G) and synaptic regulation (H). Plotted as Log2 FC at each age relative to P120. * FDR < 0.05 by DESeq2 with Benjamini-Hochberg’s correction when comparing P120 to each age.

To assess if there are broad changes in the transcriptomic profiles of astrocytes across development we performed principal component analysis (PCA) (Fig 2C). This showed that P7 and P14 astrocytes form distinct clusters, while P28 and P120 astrocytes cluster together. To investigate this further, we analyzed the number of differentially expressed genes (DEGs; FDR <0.05) between each age group. DEGs are classified into total genes (all changes detected, FPKM >1), genes that are expressed by astrocytes (IP/input >0.75), and genes that are enriched in astrocytes (IP/input >3; Fig 2D, for definitions see also (Boisvert et al., 2018)). The largest number of DEGs is between P7 and P28 (~6000 total genes), and smallest numbers between P14 to P28 (~1000 total genes), and P28-P120 (~2000 total genes). Analysis of astrocyte-enriched genes (IP/input >3) showed that ~60% of all astrocyte-enriched genes are significantly changed from P7 to P14, while only ~20% are changing between P28 to P120 (Fig 2E). This shows that most changes in astrocyte gene expression are occurring between the first and second postnatal weeks, a time of transition from synapse formation to synapse maturation, and from spontaneous to visually-evoked neuronal activity.

To determine the different astrocyte functions at each age we performed gene ontology (GO) analysis, focusing on Biological Processes terms (BP) (Fig S2F-H). This showed that genes common to all ages are enriched in GO terms related to cholesterol processing and serine synthesis, confirming the previously established important role of astrocytes in regulating brain cholesterol (Orth and Bellosta, 2012)(Fig S2G). Analysis of GO terms unique to each age showed that at P7 astrocyte genes are enriched in GO terms related to cortical development, while at P14 astrocyte genes are enriched in GO terms related to Wnt and BMP signaling pathways. Conversely, adult astrocytes (P120) are enriched in terms related to regulation of extracellular matrix assembly and contact inhibition (Fig S2H). In all we found 547 GO terms common to all ages (more than 50% of all terms identified for each age), while terms unique to each age consisted less than 10% of all terms (Fig S2F). These results suggest that, for the most part, astrocytes perform core functions that are occurring at all developmental stages, while several age-specific functions turn on and off depending on the developmental stage.

We next used this dataset to identify potential astrocyte marker genes across ages, as well as analyzing the developmental expression changes of key astrocyte genes. We determined that Apoe and Cst3 are the most highly expressed astrocyte genes at all ages (Fig 2F), while Lars2 is the most astrocyte-specific gene at all ages (Fig S2E). The expression of the known astrocyte marker Aldh1l1 is stable across all ages, making it an optimal marker for astrocytes at any age. On the other hand, S100β expression is upregulated later in development, making it a more suitable marker for adult astrocytes (Fig 2G), while vimentin expression is high at early time points, making it a good early-stage astrocyte marker. For genes that encode proteins important for astrocyte function, we found that the metabotropic glutamate receptor mGluR5 (Grm5) is most highly expressed at P7 and then declines with maturation, while the glutamate transporter Glt-1 (Slc1a2) and the connexins (Gja1, Gjb6) are significantly upregulated from P14 onwards (Fig 2G).

Finally we focused on astrocyte genes that encode proteins known to regulate neuronal synapse number, function, and maturation (Fig 2H). These include astrocyte-secreted thrombospondins (Thbs), which induce silent synapse formation. The family members expressed by VC astrocytes show divergent expression, with Thbs1 being significantly higher at P7 than later ages, whereas Thbs4 is significantly lower at P7 than P120 (Fig 2H). This temporal expression profile fits with previous studies that have demonstrated important roles for Thbs1 in initial synapse formation at P7 (Christopherson et al., 2005), and suggested roles for Thbs4 in the adult brain (Benner et al., 2013). Similarly, glypican (Gpc) family members have a divergent expression. Gpc4, which induces formation of immature synapses (Allen et al., 2012), is most highly expressed at P7 and gradually declines with maturation, whereas Gpc6 peaks between P14-P28. Gpc5, a glypican family member with yet unknown function, has low expression at P7 and is significantly increased at all later ages. Astrocyte-secreted chordin like 1 (Chrdl1) regulates synapse maturation and its expression peaks at P14 (Blanco-Suarez et al., 2018) (Fig 2H). These changing temporal expression profiles are not limited to factors that promote synapse formation. Astrocyte phagocytic receptors involved in synapse elimination, Megf10 and Mertk (Chung et al., 2013), significantly increase in expression between P7 and P14, coincident with the initiation of synapse elimination. C4b, a component of the complement cascade involved in synapse elimination is significantly upregulated at P120 compared to all younger ages (Fig 2H).

In summary, the transcriptomic analysis reveals significant changes in astrocyte gene expression across development, with the most prominent changes being between P7 and later ages, a time between synapse initiation and maturation. It further points out the differential developmental expression patterns of synapse-regulating genes, such as Gpc4 peaking during synapse initiation, and Chrdl1 peaking after eye opening, when synapses begin to mature. These data can be further utilized to infer functional changes in astrocytic roles and inform further studies on astrocyte development. The complete dataset and GO term list are available in Tables S2A-B.

### Synapse-regulating genes in astrocytes show differential spatio-temporal expression

Given the layer-specific alterations in astrocyte and synapse number that occur over development (Fig 1), along with developmental changes in astrocyte synapse-regulating gene expression shown by bulk RNA sequencing (Fig 2), we next asked if there are correlated layer-specific changes in astrocyte synapse-regulating genes that could be contributing to these effects. For example, the largest increase in GluA2 occurs in upper cortical layers between P7 and P14 (Fig 1M), so we hypothesized that astrocyte Chrdl1 expression would follow the same spatial pattern. To determine layer-specific developmental changes in mRNA expression of synapse-regulating genes we performed single-molecule fluorescent in situ hybridization (smFISH; RNAscope) on brain sections of Aldh1l1-GFP mice and probed for 7 genes that regulate distinct aspects of synaptogenesis: active synapse-regulating - glypicans (Gpc) 4, 5, 6; synapse maturation regulating – chordin-like 1 (Chrdl1); and silent synapse-regulating – thrombospondins (Thbs) 1, 2, 4. Expression of each gene was analyzed within the territory of GFP positive astrocytes in each cortical layer and at 4 developmental time points: P4, P7, P14 and P28, when the most alterations in astrocyte and synapse development occur (Fig 1-2, Fig 3A-B, S3A). A negative control probe was used to determine the minimal signal threshold of detection (Fig S3H-I).

**Figure 3.**
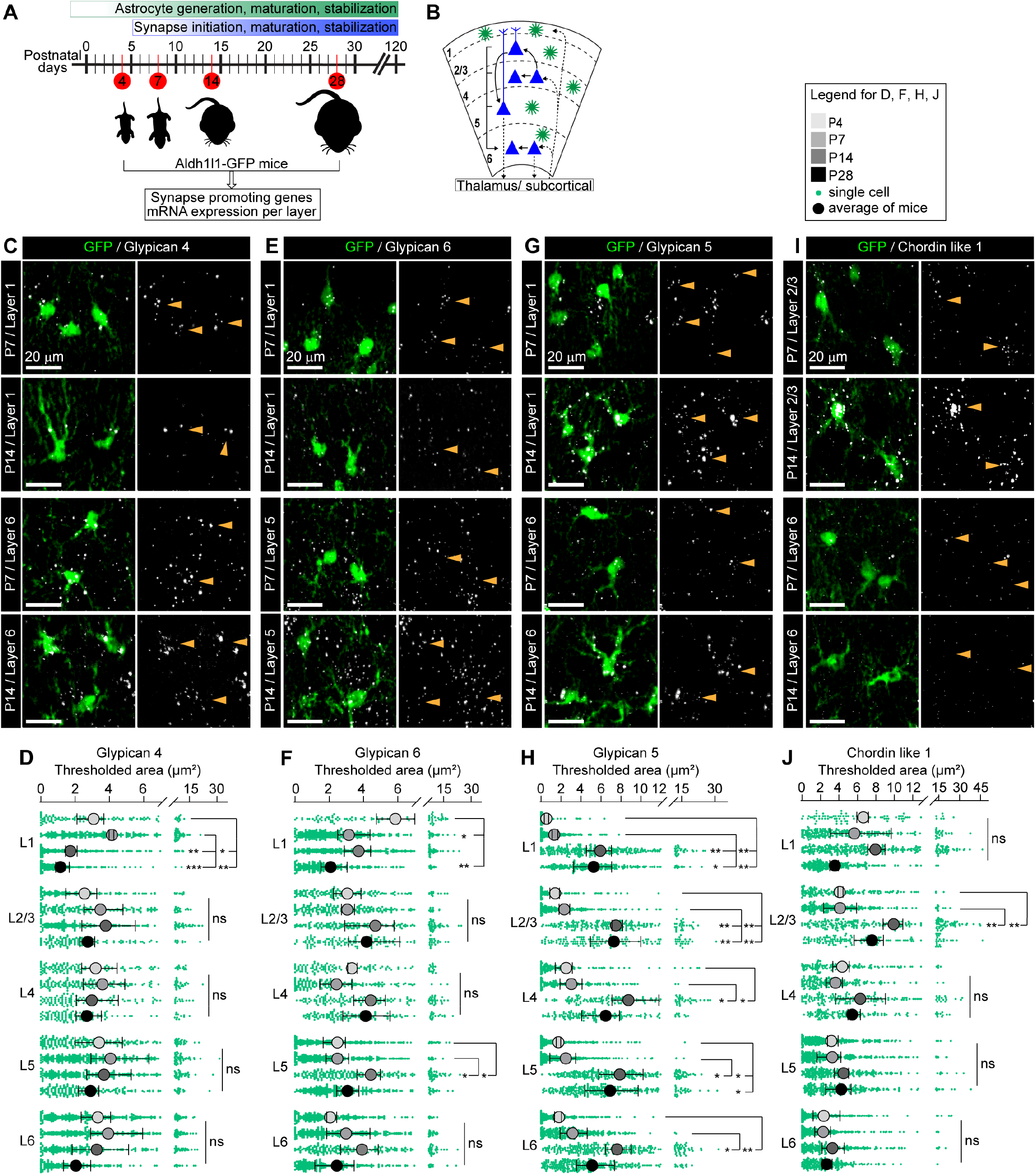
Synapse-regulating genes in astrocytes show differential spatio-temporal expression. See also Fig S3, Table S3. **A.** VCs were collected from Aldh1l1-GFP mice at different post-natal ages corresponding to synapse development. In situ hybridization (ISH) was performed to assess mRNA level for synapse-regulating genes in astrocytes in each layer. **B.** Diagram of visual cortex depicting neuronal (blue) laminar arrangement and connectivity (arrows). Astrocytes (green) present in all layers of the VC. **C, E, G, I.** Example images showing Gpc4, Gpc6, Gpc5 or Chrdl1 mRNA (white) in astrocytes (green) at each age and layer as labeled. Merged panel on the left, single-channel probe panel on the right. Arrowheads in single-channel panel mark astrocyte cells on the left. Scale bars = 20 μm. **D, F, H, J.** Quantification of C, E, G, I respectively. **D.** Gpc4 expression is reduced at P14 specifically in L1. **F.** Gpc6 expression is increased at P14 in L5. **H.** Gpc5 expression is increased at P14 in all layers. **J.** Chrdl1 expression is increased at P14 in L2/3. Data presented as scatter with mean + range, large circles average calculated from data per mouse and colored according to time point, green dots signal per astrocyte. N=3 mice/age, n=~50-350 astrocytes/per age; averages and statistical analysis are calculated based on N=3 i.e. data per mouse. *P ≤ 0.05 **P<0.01, ***P<0.001, ns (not significant) by one-way ANOVA comparing expression between time points within each layer.

We first analyzed glypicans, factors that promote active synapse formation. Bulk RNAseq showed that astrocyte Gpc4 expression is highest at P7 and gradually declines with maturation. Layer-specific analysis, however, shows that these differences are driven by astrocytes in L1: Gpc4 expression decreases between P7 and P14 only in L1 astrocytes, staying stable across development in all other layers (Thresh area (μm^2^) L1: P4 3.08 ±0.49; P7 4.14 ±0.08; P14 1.72 ±0.2; P28 1.14 ±0.26; Fig 3C,D, Table S3A,B). Gpc6, on the other hand, peaks at P14-P28 in bulk sequencing, and layer-specific analysis showed that this increase occurs in the majority of astrocytes, increasing between P7-P14 in L2-5 (with significant upregulation in L5), and remaining high at P28 (Thresh area (μm^2^) L5: P4 2.48 ±0.42; P7 2.48 ±0.38; P14 4.42 ±0.41; P28 3.06 ±0.38; Fig 3E,F, Table S3A,B). Gpc5 is strongly upregulated at P14 in astrocytes in all layers and remains high at P28, matching the bulk RNAseq data (Thresh area (μm^2^) L1: P4 0.53 ±0.13; P7 1.33 ± 0.18; P14 5.93 ±0.73; P28 5.27 ±1.11; Fig 3G,H, Table S3A,B).

The expression of the synapse maturation factor Chrdl1 peaks at P14 in the bulk RNAseq data (Fig 2G). Spatial analysis revealed that this increase from P7 to P14 is layer-specific, with the largest increase in Chrdl1 occurring in upper layer astrocytes with significant upregulation in L2/3 (Thresh area (μm^2^) L2/3: P4 4.1 ±0.18; P7 4.11 ±1.03; P14 9.93 ±0.62; P28 7.59 ±0.98; Fig 3I,J, Table S3A,B). Further, at its peak of expression, Chrdl1 is highest in L1-4 astrocytes compared to L5-6, demonstrating a heterogeneous expression across layers (Fig 3I,J, S3A, Table S3A,B). Finally, we analyzed astrocyte thrombospondins, factors that induce silent synapse formation (Fig S3B-G). Thbs mRNA levels in the VC are much lower at their peak expression than glypicans or Chrdl1, consistent with our bulk RNAseq results (Fig 2G, Table S2A), and previous studies showing low thrombospondin expression in the resting state and an upregulation by learning or injury (Nagai et al., 2019; Tyzack et al., 2014). The only significant developmental changes observed in our experiments are an increase in Thbs2 in L4-5 at P28 (Thresh area (μm^2^) L5: P4 0.65 ±0.1; P7 0.7 ±0.19; P14 0.98 ±0.11; P28 1.77 ±0.27; Fig S3D,E, Table S3A,B), and an increase in Thbs4 in astrocytes in all layers at P28, again consistent with the bulk sequencing (Thresh area (μm^2^) L1: P4 0.05 ±0.01; P7 0.13 ± 0.06; P14 0.3 ±0.08; P28 1.93 ± 0.49; Fig S3 F,G, Table S3A,B).

In all, we have determined the spatio-temporal expression profile of key astrocyte synapse-regulating factors, identifying divergent developmental and layer-specific expression patterns within the same families of genes. These findings strongly suggest that astrocyte expression of synapse-regulating genes is closely tied to the developmental stage of the cortex, which features both changes in neuronal and astrocyte activities across development.

### Neuronal activity tunes astrocyte expression of synapse-regulating genes

Having found broad differences in astrocyte expression of synapse-regulating genes across cortical layers, we next asked what regulates these layer-specific changes between P7 and P14. Given that during this time mouse eye-opening occurs, we hypothesized that blocking visually-evoked neuronal activity would disrupt these changes. Previous work, as well as our *in vitro* experiments using cultured astrocytes and neurons, show that Gpc4 mRNA expression (Hasel et al., 2017) and protein secretion from astrocytes (Fig S4A) are significantly reduced in the presence of neurons, suggesting neurons can influence expression and release of synapse-regulating factors from astrocytes.

To prevent glutamate release from thalamic neurons that innervate the VC and relay information from the retina, we knocked out the vesicular glutamate transporter VGlut2 from neurons in the dLGN. Knockout of VGlut2 has been previously shown to abolish presynaptic release of glutamate in VGlut2 expressing neurons, in full or conditional knockout mouse models (Wallén-Mackenzie et al., 2010). We crossed VGlut2^f/f^ (labeled as VGlut2 WT) mice to an RORα cre line (VGlut2^f/f;cre^ labeled as VGlut2 cKO), where cre recombinase is expressed in neurons in the thalamus including the dLGN (Fig 4A) (Chou et al., 2013; Farhy-Tselnicker et al., 2017). Immunostaining experiments showed a significant decrease in VGlut2 signal in the VC of VGlut2 cKO mice compared to WT at P14 (Fig 4B), with no overall effect on VGlut1 which marks cortico-cortical terminals (Fig 4C). A lack of VGlut2 signal in the VC could also result from an absence of thalamic axons innervating their target regions. To test whether that is the case we crossed RORα cre and VGlut2 cKO mice with an Ai14 tdTomato reporter line to visualize dLGN axons (Fig 4A). All cre positive mice (WT, VGlut2 cHet, and VGlut2 cKO) showed a comparable number and volume of tdTomato labeled projections in L1 and L4 of the VC (Fig 4A, S4B-D). Analysis of VGlut2 puncta colocalized with tdTomato positive axons showed a significant decrease in number in VGlut2 cHet and VGlut2 cKO compared to WT (Fig S4B, D). These results show that in the VGlut2 cKO mice thalamic axons are present at their target layers in the VC but lack VGlut2 (as has been shown in studies performing similar manipulations (Li et al., 2013; Zechel et al., 2016)), suggesting they are functionally silent

**Figure 4.**
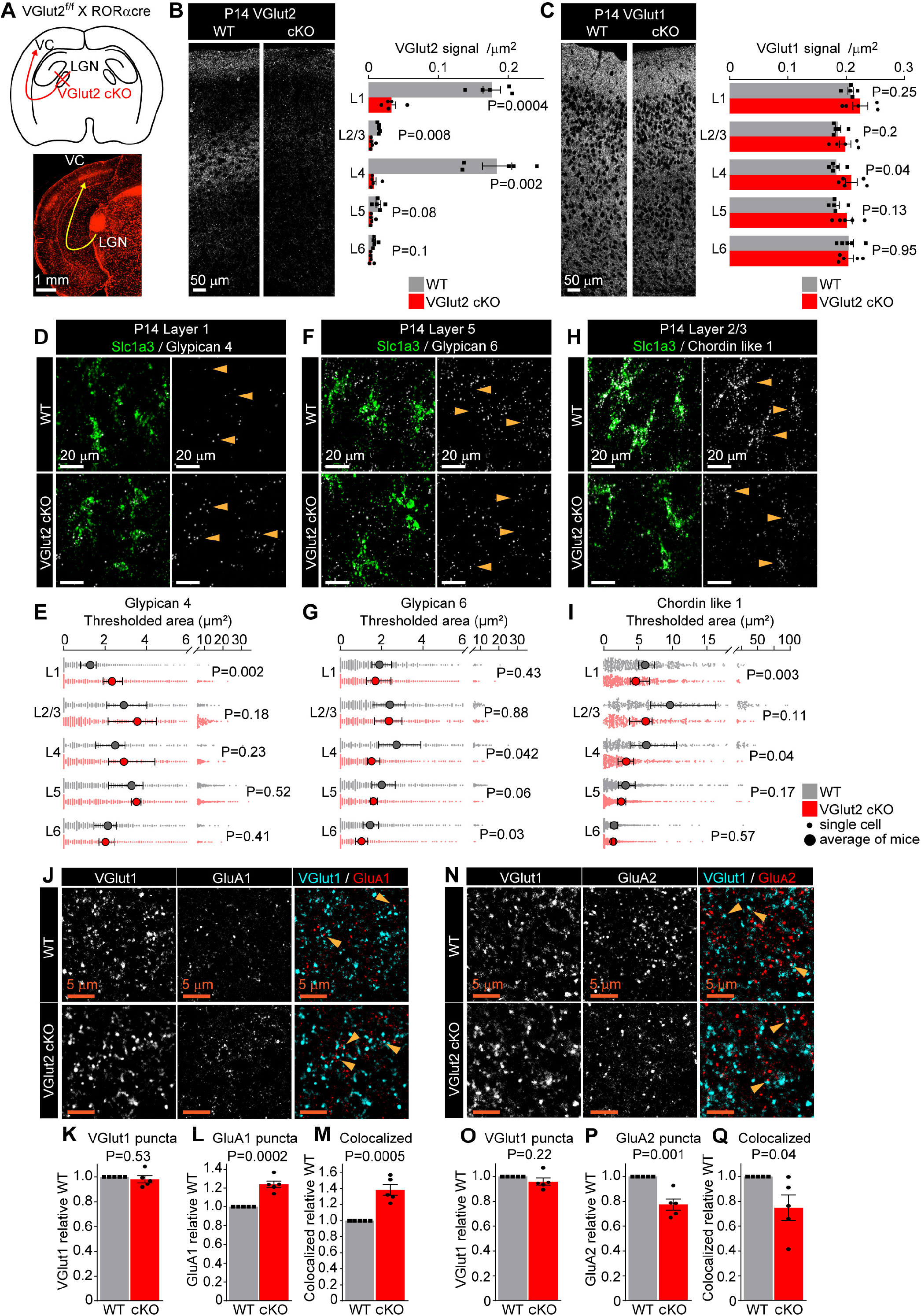
Neuronal activity tunes astrocyte expression of synapse-regulating genes. See also Fig S4, Table S4. **A.** Schematic of the experiment: VGlut2 is removed from presynaptic terminals of neurons in the lateral geniculate nucleus of the thalamus (LGN), that project to the visual cortex (VC), by crossing VGlut2 f/f mouse (WT) with RORαcre mouse line (VGlut2 cKO). Bottom: image of tdTomato reporter expression in the LGN and the VC, when RORαcre mouse is crossed with cre-dependent tdTomato reporter mouse. **B.** VGlut2 expression in the VC is significantly reduced in VGlut2 cKO mice. Example images of VGlut2 immunostaining in each genotype and quantification of the thresholded signal within each cortical layer. **C.** VGlut1 level is unaltered in VGlut2 cKO mice. Example images of VGlut1 immunostaining and quantification. In B, C. plots show mean signal ± s.e.m. Squares and circles are the average of signal in each mouse. N=5 mice/genotype. Scale bar = 50 μm. Statistical analysis by t-test within each layer. P-value on each plot. **D-I.** mRNA expression of astrocyte synapse-regulating genes is altered in VGlut2 cKO at P14. **D, F, H.** Example images of in situ hybridization of Gpc4, Gpc6 and Chrdl1 mRNA (white) as labeled; astrocyte marker Slc1a3 (Glast, green). Merged panel on the left, single-channel probe panel on the right. Arrowheads in single-channel panel mark astrocytes. Scale bar = 20 μm. **E, G, I.** Quantification of D, F, G respectively. **E.** Gpc4 mRNA expression is increased in L1; **G.** Gpc6 mRNA expression is decreased in L4-6; **I.** Chrdl1 mRNA expression is decreased in L1-4 in VGlut2 cKO mice. Data presented as scatter with mean + range, large circles are the average signal calculated from data per mouse. Grey or red dots are signals in individual astrocytes in WT and VGlut2 cKO respectively. N=5 mice/genotype, n=~200-450 astrocytes/ per age total (average and statistical analysis is calculated based on N=5 i.e. per mouse). Statistical analysis by paired t-test within each layer. P value on each plot. **J-M.** Increase in GluA1 protein levels and colocalization between GluA1 and VGlut1 in L1 of the VC in VGlut2 cKO mice at P14. **J.** Example images from WT (top) and cKO (bottom), VGlut1 in cyan and GluA1 in red. **K, L, M.** Quantification of VGlut1, GluA1 and colocalized puncta respectively, normalized to WT. **N-Q.** Decrease in GluA2 protein levels and colocalization between GluA2 and VGlut1 in L1 of the VC in VGlut2 cKO mice at P14. **N.** Example images from WT (top) and cKO (bottom), VGlut1 in cyan and GluA2 in red. **O, P, Q.** Quantification of VGlut1, GluA2 and colocalized puncta respectively, normalized to WT. In K-M and O-Q data presented as mean ± s.e.m, squares and circles each mouse. N=5 mice/ genotype. Arrowheads mark representative colocalized puncta in J, N. Scale bar = 5 μm. Statistical analysis by t-test, p-value on each plot.

Does the lack of visually-evoked neuronal activity impact the expression of synapse-regulating genes in astrocytes? To address this we performed smFISH at P14 probing for Gpc4, Gpc6 or Chrdl1, genes which showed the most robust layer-specific developmental changes in expression between P7 and P14, along with a probe for the glutamate transporter Glast (Slc1a3)(Ullensvang et al., 1997) to label astrocytes. The number of cortical astrocytes marked by Glast, and the expression level of Glast mRNA, are not affected by VGlut2 cKO in any cortical layer at P14, showing that gross astrocyte development proceeds normally in the absence of visual input (Fig S4E,F). During normal development Gpc4 expression significantly decreases between P7 and P14 specifically in L1 astrocytes (Fig 3C,D). In VGlut2 cKO mice this change no longer occurs (Fig 4D,E). Gpc4 expression is significantly increased in VGlut2 cKO compared to WT at P14 specifically in L1 astrocytes, and unchanged in all other layers (Thresh area (μm^2^): L1 WT 1.27 ± 0.13; cKO 2.32 ± 0.15; Fig 4D,E, Table S4A,B). Gpc6 expression is normally increasing in astrocytes in deep layers between P7 and P14 (Fig 3E,F). In the absence of thalamic glutamate release, Gpc6 is lower in astrocytes in layers 4 and 5 than in the WT at P14, showing that the normal developmental upregulation has been blocked (Thresh area (μm^2^): L4 WT 2.73 ± 0.36; cKO 1.53 ± 0.11; Fig 4F,G, Table S4A,B). Similarly, Chrdl1 expression normally increases in upper layer astrocytes between P7 and P14 (Fig 3I,J), however it is significantly decreased in VGlut2 cKO compared to WT specifically in astrocytes in upper layers (1-4), and not affected in deep layers at P14 (Thresh area (μm^2^): L1 WT 5.98 ± 0.41; cKO 4.62 ± 0.51; Fig4H,I, Table S4A,B). To determine if these alterations are due to the loss of visually-evoked neuronal activity at P14, rather than due to sustained suppression of glutamate release from thalamic neurons throughout development, we also analyzed the expression of Gpc4, Gpc6 and Chrdl1 at P7, and observed no difference in expression of any of these genes in the cKO (Fig S4I-K), nor any difference in the number of cortical astrocytes marked by Glast, or the expression level of Glast mRNA (Fig S4G,H). These results show that during visual cortex development from P7 to P14, glutamate release from thalamic neurons regulates the developmental expression of astrocyte Gpc4, Gpc6 and Chrdl1 in a layer-specific manner.

What is the consequence of altered expression of astrocyte synapse-regulating genes on synaptic development? To address this we performed immunostaining for pre- and postsynaptic proteins in VGlut2 cKO mice and their WT controls in L1 of VC at P14. We first analyzed cortico-cortical synapses marked by VGlut1 and found that, as in the low-resolution characterization (Fig 4C), there is no change in VGlut1 puncta in the absence of VGlut2 (Fig 4J,K,N,O). In the case of GluA1 containing AMPARs, which are regulated by Gpc4, we found a significant increase in the number of GluA1 puncta and their colocalization with VGlut1 in the VGlut2 cKO, correlating with the observed increase in Gpc4 (GluA1 FC 1.24 ±0.04; Fig 4J,L; Coloc FC 1.38 ±0.07 Fig 4J,M). For GluA2 containing AMPARs, which are regulated by Chrdl1, we found a significant decrease in both the number of GluA2 puncta and the number of colocalized GluA2-VGlut1 puncta in VGlut2 cKO mice compared to WT, correlating with the observed decrease in Chrdl1 (GluA2 FC 0.77 ±0.04 Fig 4N,P; Coloc FC 0.75 ±0.1 Fig 4 N,Q). We asked if similar effects are also present at thalamo-cortical synapses. Since VGlut2 is absent in cKO mice we used VGlut2f/f; RORαcre;tdTomato to label thalamic axons (Fig 4A, Fig S4B), and identified presynaptic active zones within tdTomato axons by immunostaining for the pre-synaptic marker bassoon (Fig S4L,P). We found no difference in the number of bassoon puncta colocalized with tdTomato between the WT and cKO mice, a further indication that synapses form in the absence of VGlut2 (Fig S4L-M, P-Q), and fitting with findings from mice that globally lack presynaptic release (Verhage et al., 2000). As is the case for cortico-cortical synapses, we found an increase in GluA1 and colocalization of GluA1 with bassoon and tdTomato (GluA1 FC 1.21 ±0.09 Fig S4L,N; Coloc FC 1.36 ±0.23 Fig S4L,O), and a decrease in total GluA2 and GluA2-bassoon synapses (GluA2 FC 0.70 ±0.08 Fig S4P,R; Coloc FC 0.65 ±0.13 Fig S4P,S). These results show that synaptic GluA1 and GluA2 levels are altered in VGlut2 cKO VC in the direction which follows the change in astrocytic expression of Gpc4 (which recruits GluA1) and Chrdl1 (which recruits GluA2) in L1. These correlated changes in astrocyte genes and synaptic proteins suggest a delay in synapse maturation in the VC at P14 in the absence of thalamo-cortical glutamate release.

### Astrocyte calcium signaling regulates expression of synapse-regulating genes

Since we observed that changes in neuronal activity can regulate expression of astrocyte synapse-regulating genes, we next asked whether perturbing the astrocyte response to neuronal activity affects the expression of Gpc4, Gpc6 and Chrdl1. Astrocytes express many neurotransmitter receptors, in particular GPCRs, and respond to the majority of neurotransmitters with increased intracellular calcium (Kofuji and Araque, 2020; Porter and McCarthy, 1997). In the case of somal increases in calcium, which have the potential to regulate expression of activity-regulated genes, most of this increase is mediated by the release of calcium from intracellular stores via Ip3r2 (Itpr2) (Srinivasan et al., 2015). We therefore asked if blunting astrocyte calcium signaling by removing store-mediated calcium release using Ip3r2 KO mice (Fig 5A,B) has an impact on the expression of synapse-regulating genes.

**Figure 5.**
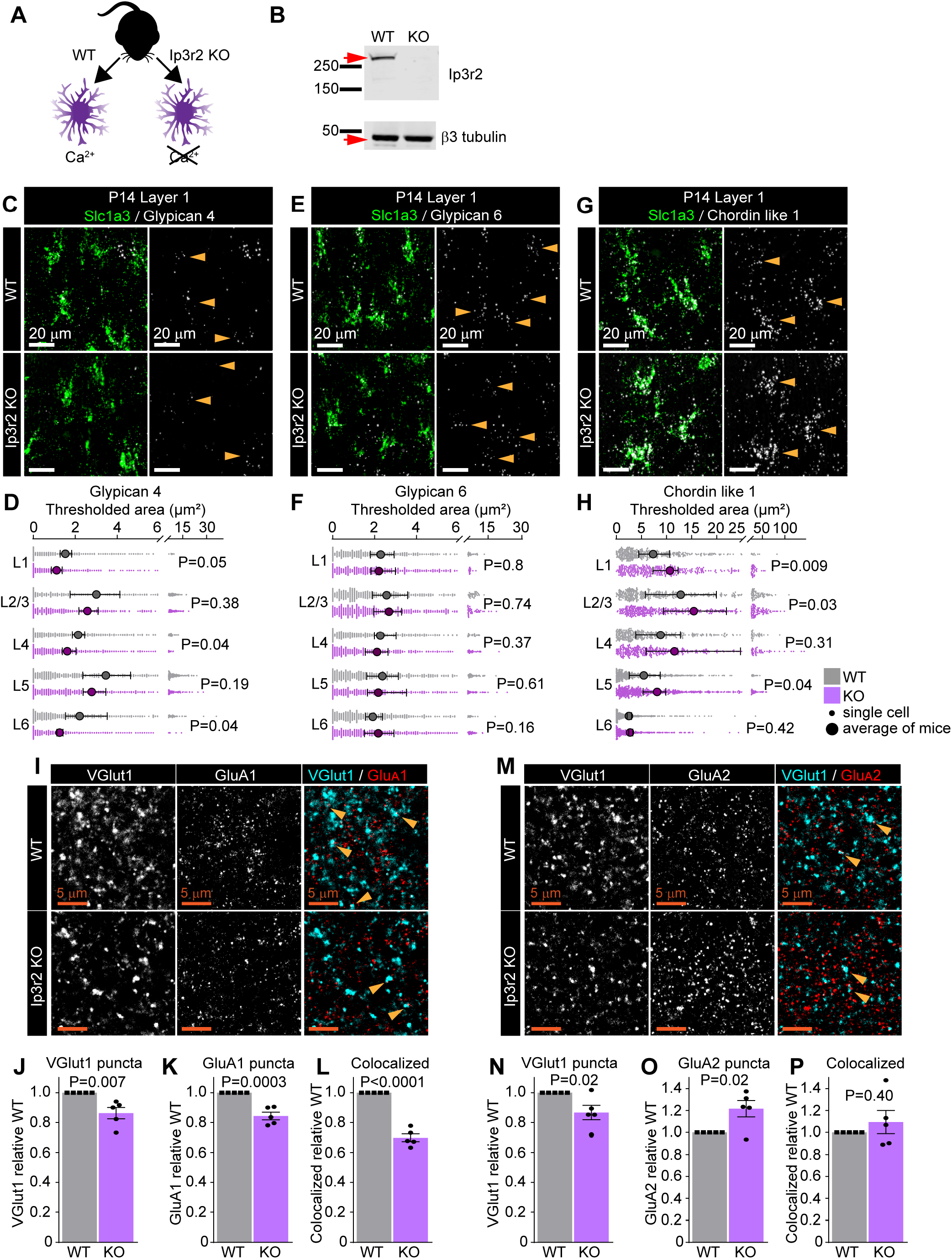
Astrocyte calcium signaling regulates expression of synapse-regulating genes. See also Fig S5, Table S4. **A.** Schematic of comparison. Lack of the Ip3r2 receptor results in diminished Ca^2+^ transients in astrocytes. **B.** Validation of Ip3r2 KO model. Western blot shows absence of Ip3r2 signal in VC of KO mice. **C-H.** mRNA expression of astrocyte synapse-regulating genes is altered in Ip3r2 KO mice at P14. **C, E, G.** Example images of in situ hybridization of Gpc4, Gpc6 and Chrdl1 mRNA (white); astrocyte marker Slc1a3 (Glast, green). Merged panel on the left, single-channel probe panel on the right. Arrowheads in single-channel panel mark astrocytes. Scale bar = 20 μm. **D, F, H.** Quantification of C, E, G respectively. **D.** Gpc4 mRNA expression is decreased in several layers of the VC in Ip3r2 KO mice. **F.** Gpc6 mRNA expression is unaltered in the VC in Ip3r2 KO mice. **H.** Chrdl1 mRNA expression is increased in several layers of the VC in Ip3r2 KO mice. Data presented as scatter with mean + range, large circles are calculated from average signal per mouse. Grey or purple dots are signal per astrocyte in WT and Ip3r2 KO respectively. N=5 mice/genotype, n=~200-450 astrocytes/ per age total (average and statistical analysis is calculated based on N=5 i.e. per mouse). Statistical analysis by paired t-test within each layer. P value on each plot. **I-L.** Decrease in VGlut1, GluA1 protein levels, and colocalization between GluA1 and VGlut1 in L1 of the VC in Ip3r2 KO mice at P14. **I.** Example images from WT (top) and KO (bottom), VGlut1 in cyan and GluA1 in red. **J, K, L.** Quantification of VGlut1, GluA1 and colocalized puncta respectively, normalized to WT. **M-P.** Decrease in VGlut1, and increase GluA2 protein levels, with no change in colocalization between GluA2 and VGlut1 in L1 of the VC in Ip3r2 KO mice at P14. **M.** Example images from WT (top) and KO (bottom), VGlut1 in cyan and GluA2 in red. **N, O, P.** Quantification of VGlut1, GluA2 and colocalized puncta respectively, normalized to WT. In J-L and N-P data presented as mean ± s.e.m, squares and circles represent mice. N=5 mice/genotype. Arrowheads mark representative colocalized puncta. Scale bar = 5 μm. Statistical analysis by t-test, p-value on each plot.

To determine this we performed smFISH on the VC of P14 Ip3r2 KO and WT mice, marking astrocytes with a probe against Glast along with Gpc4, Gpc6, or Chrdl1. At P14 knocking out Ip3r2 does not affect the number of astrocytes or the expression levels of Glast (Slc1a3), showing astrocytes develop grossly normally when store-mediated calcium release is diminished (Fig S5A,B). However, loss of Ip3r2 does impact expression of synapse-regulating genes. In the case of Gpc4, the mRNA level is reduced in astrocytes in all layers, with a significant decrease occurring in L1, 4 and 6 (Thresh area (μm^2^): L1 WT 1.53 ± 0.13; KO 1.11 ± 0.1; Fig 5C,D, Table S4A,B). For Gpc6 there is no difference in the mRNA level between Ip3r2 KO and WT in astrocytes in any layer (Fig 5E,F, Table S4A,B). Chrdl1 expression is increased in astrocytes in all layers, with a significant increase occurring in L1, 2/3 and 5 (Thresh area (μm^2^): L1 WT 7.32 ± 1.09; KO 10.64 ± 0.97; Fig 5G,H, Table S4A,B). To ask if these alterations are present throughout development we performed the same analysis at P7. As with P14, at P7 there is no change in astrocyte number or Glast mRNA signal (Fig S5C,D). In the case of Gpc4 and Chrdl1 there is no difference in expression between Ip3r2 KO and WT at P7 (Fig S5E,G), whereas for Gpc6 there is a significant increase in the Ip3r2 KO restricted to L4 (Fig S5F). Therefore, in contrast to the layer-specific alterations in gene expression in the VGlut2 cKO mice, the effects of removing Ip3r2 are impacting astrocytes in all layers and do not strictly follow the developmental trajectory. This suggests a broad requirement for astrocyte calcium signaling in all astrocytes to maintain the correct level of gene expression, and that the signals to do this come from multiple sources and are not restricted to thalamic inputs.

What are the consequences of diminished astrocyte calcium signaling and altered expression of synapse-regulating genes on synaptic development? As with the VGlut2 cKO, we addressed this by performing immunostaining for presynaptic terminals (VGlut1 or VGlut2) and postsynaptic AMPAR subunits (GluA1 or GluA2) in Ip3r2 KO mice and WT controls in L1 of VC at P14. We found that the numbers of both cortico-cortical presynaptic terminals marked by VGlut1 and thalamo-cortical presynaptic terminals marked by VGlut2 are significantly decreased in Ip3r2 KO mice compared to WT, demonstrating a global deficit in synapse formation in the absence of astrocyte calcium signaling (VGlut1 from GluA1: FC 0.86 ±0.04 Fig 5I,J; VGlut1 from GluA2: FC 0.87 ±0.05 Fig 5M,N. VGlut2 from GluA1: FC 0.89 ±0.05 Fig S5H,I; VGlut2 from GluA2: FC 0.80 ±0.05 Fig S5L,M). For GluA1 containing AMPARs, which are regulated by Gpc4, we found a significant decrease in the total number of puncta and their colocalization with VGlut1 and VGlut2 in the Ip3r2 KO, correlating with the observed decrease in Gpc4 mRNA (VGlut1-GluA1: GluA1 FC 0.84 ±0.03 Fig 5I,K; Coloc FC 0.70 ±0.03 Fig 5I,L. VGlut2-GluA1: GluA1 FC 0.83 ±0.03 Fig S5H,J; Coloc FC 0.74 ±0.03 Fig S5H,K). For GluA2 containing AMPARs, which are regulated by Chrdl1, we found a significant increase in their number, correlating with the observed increase in Chrdl1 (VGlut1-GluA2: GluA2 FC 1.22 ±0.07 Fig 5M,O. VGlut2-GluA2: GluA2 FC 1.14 ±0.03 Fig S5L,N). The number of colocalized presynaptic puncta and GluA2 is, however, unchanged, likely due to the opposing decrease in presynaptic puncta and increase in GluA2 (VGlut1-GluA2 Coloc FC 1.09 ±0.1 Fig 5M,P; VGlut2-GluA2 Coloc FC 1.00 ±0.08 Fig S5L,O).

Taken together these results show that GluA1 and GluA2 levels are altered in Ip3r2 KO VC in the direction which follows the change in astrocytic expression of Gpc4 (which recruits GluA1) and Chrdl1 (which recruits GluA2). This strongly suggests that both astrocytes and neurons play an important role in regulating the expression of synapse-regulating genes, and subsequently AMPAR subunit protein levels, and the final expression levels arise from the complex interaction between these two cell types.

### Unbiased determination of astrocyte diversity and activity-regulated genes in the developing visual cortex

Having found that multiple synapse-regulating genes in astrocytes show layer-specific enrichment, and that these patterns are regulated by neuronal and astrocyte activity, we next asked if these findings are specific to synapse development, or if other astrocyte genes show a similar pattern. To address this using an unbiased approach we performed single-nucleus RNA sequencing of glial cells isolated from the P14 VC of wild type, VGlut2 cKO, and Ip3r2 KO mice. To isolate the glial cell populations we immunostained VC nuclei in suspension with an antibody against the neuronal marker NeuN, and performed FACS to select the NeuN-negative population (Fig 6A). We used the Chromium 10X system to isolate individual glial nuclei and performed RNA sequencing to quantify mRNA levels (Fig 6A). This identified 22,781 cells in the VGlut2 condition (cKO and WT) (Fig 6B), and 21,240 cells in the Ip3r2 condition (KO and WT) (Fig S6A). Initial clustering analysis determined 17 distinct cell populations in both models, with the majority of cells detected clustered within the main glial cell types: astrocytes, microglia, and oligodendrocyte lineage cells (Fig 6B,C, S6A,B). Just two clusters enriched for neuronal markers are present, showing that the NeuN depletion had been successful.

**Figure 6.**
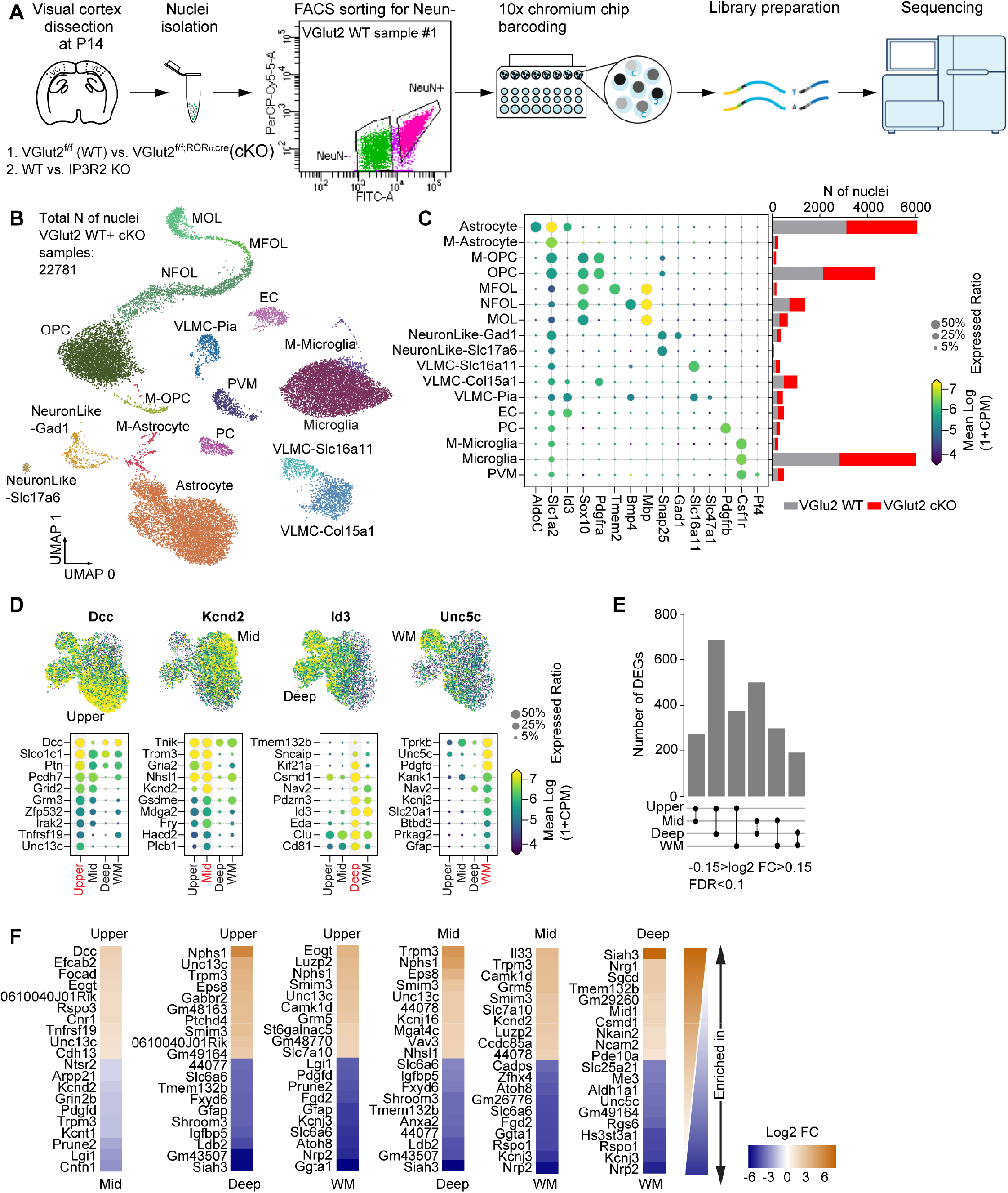
Unbiased determination of astrocyte layer-enriched genes. See also Fig S6, Table S5. **A.** Outline of experiment: VCs collected from VGlut2 cKO, Ip3r2 KO and their respective WT controls at P14. Nuclei were isolated from VCs, and sorted for NeuN negative population (glia) using flow cytometry. Sorted nuclei were loaded onto 10x Chromium chip, each nucleus barcoded, followed by library preparation and sequencing. N=8 samples total. **B.** UMAP clustering of different cell types identified in the NeuN negative population of the combined samples from VGlut2 WT and cKO mice. 17 clusters were identified including the 3 main types of glia: astrocytes, oligodendrocytes and microglia, as well as endothelial cells, and two sub-types of neurons (Abbreviations are: M-astrocyte – mitotic astrocyte; M-OPC – mitotic oligodendrocyte precursor cell; OPC – oligodendrocyte precursor cell; MFOL - myelin-forming oligodendrocyte; NFOL - newly formed oligodendrocyte; MOL - mature oligodendrocyte; VLMC-vascular and leptomeningeal cell; EC – Endothelial cell; PC - pericyte; PVM - perivascular macrophage). **C.** Expression level of select marker genes for each cluster. Circle size denotes expression ratio (percent cells expressing the gene), color represents expression level (in Log2 CPM). Bar chart on the right is the number of cells identified for each genotype as labeled. Similar cell numbers were identified for VGlut2 WT and cKO groups for each cluster. **D.** Unbiased clustering analysis identified 4 subpopulations of astrocytes in the P14 VC. Populations annotated to Upper, Mid, Deep and White matter types following comparison with published datasets. Top panel shows the expression level of select marker genes that label a particular population as indicated. Each dot represents a single nucleus, color represents expression level in Log2 CPM. Bottom panels show a select list of 10 genes that are highly expressed in each population as indicated. Size of the circle is expression ratio; color is expression level (log2 CPM). **E.** Pairwise comparison identified ~200-700 DEGs between astrocyte populations from WT mice from the VGlut2 analysis. Criteria for DEG selection: Log2 FC between −0.15 and 0.15; FDR <0.1. **F.** Heatmap showing top 20 DEGs from each pairwise comparison (**E**) showing genes enriched in one population vs the other (top vs bottom labels). Colors represent Log2 Fold change (FC) between each population.

### Determination of astrocyte layer-enriched genes in the developing wild-type VC

We focused our downstream analysis on astrocytes. A second round of unbiased clustering of the astrocyte population identified 4 distinct groups (Fig 6D, S6D). By comparing the genes enriched in each cluster with datasets in the literature, we determined these to anatomically correspond to upper (up) (L1-2/3), middle (mid) (L2/3-5), deep (L5-6) layer, and white matter (WM) astrocytes (Fig S6C) (Batiuk et al., 2020; Bayraktar et al., 2020; Lanjakornsiripan et al., 2018). For example, Id3 is enriched in deep layer astrocytes and Gfap is enriched in white matter astrocytes (Fig 6D, S6D). We also determined the fractions of astrocytes present in each group and found that this corresponds to the fractions we identified via anatomical cell counts (Fig S6C,E), showing that the process of nuclear isolation has captured astrocytes in levels that reflect their *in vivo* abundance.

We first asked how many DEGs are present between the different layer groups by performing pairwise comparisons, using the WT astrocytes from either model as the input cells. This identified between 200-700 DEGs depending on the layer groups compared in the VGlut2 cKO model, with similar numbers obtained in the WT astrocytes from the Ip3r2 model, demonstrating reproducibility of the results (Fig 6E,F, S6F). The most numerous and robust differences are between the deep and upper layer astrocytes, with over 700 DEGs and up to 6-fold log2 FC in expression level (Fig 6E,F, S6F). On the other hand, upper and mid astrocytes are the most similar, with about 200 DEGs and 2-fold log2 FC maximal difference in mRNA level (Fig 6F, Table S5A,B).

Next, we asked how astrocyte marker, function, and synapse-regulating genes highlighted in the bulk RNAseq dataset (Fig 2G,H) are expressed across layers (Fig S6G). Overall, we found a positive correlation between levels of gene expression obtained by the two sequencing methods, meaning, genes that were shown to be highly expressed in the bulk dataset (such as ApoE), were also highly expressed in the snRNAseq dataset (Fig S6G). Unsurprisingly, sequencing of bulk RNA samples was more sensitive in detecting the low expressed genes, such as Gpc4, Tgfb1 and Thbs1, which were below detection level in the snRNAseq dataset. Of genes that showed detectable expression levels with the snRNAseq method, most exhibited similar expression levels across layers with some notable exceptions. For example, Gfap and Aqp4 expression is higher in deep and WM astrocytes than in upper and mid groups, while the expression of connexin 43 (Gja1) is highest in deep layer astrocytes compared to all other groups (Fig S6G).

To identify potential astrocyte layer markers we plotted the top 20 most differentially expressed genes for each pairwise comparison (Fig 6F). We found Dcc enriched in the upper layer group, consistent with previous studies (Bayraktar et al., 2020; Lanjakornsiripan et al., 2018). In addition, we found several layer-enriched genes that are absent from published datasets such as Kcnd2, a gene that is enriched in the mid-layer group, Tmem132b which is enriched in the deep astrocytes, and Unc5c, which is enriched in the white matter group. Importantly, while there are differences in astrocyte gene expression across layer groups, these are mostly gradients of gene expression, suggesting the astrocyte layer groups are on a continuum rather than distinct cell types (Fig 6D, S6D)(Bayraktar et al., 2020; John Lin et al., 2017). GO terms analysis of genes enriched in each cluster identified between 300-600 terms significantly enriched per cluster, with 140 terms that are common to all 4 clusters, and between 40-120 terms that are unique to each cluster (Fig S6H). Terms with the highest gene ratio in the upper astrocyte cluster include pathways related to signal transduction and potassium ion homeostasis, whereas mid astrocytes are enriched in terms related to GABAergic signaling and PSD95 clustering. Genes belonging to the deep astrocyte cluster are enriched in GO terms related to the regulation of pre- and post-synapse organization, while WM astrocyte genes are enriched with pathways related to axonal guidance, maintenance and signaling (Fig S6I,J, Table S5C). These results show that wild-type astrocytes in the developing VC are transcriptomically diverse, but not distinct, in accordance with previous studies in which astrocyte diversity was assessed at a similar developmental stage (Bayraktar et al., 2020).

### Global astrocyte gene expression changes following silencing of neuronal or astrocyte activity

Given we found that astrocyte synapse-regulating genes are regulated by both neuronal and astrocyte activity, we next asked what other astrocyte genes are affected by these activity manipulations. To increase the power of our analysis we combined the 4 astrocyte subpopulations into one group for each genotype and used this combined group to identify DEGs between the WT and KO. We found 61 DEGs for the VGlut2 cKO model, and 131 DEGs for the Ip3r2 model (Fig 7A-C, Table S6A). Performing the same analysis on two other abundant glial populations, OPCs and microglia, showed 28 DEGs for OPCs and 24 DEGs for microglia in the VGlut2 cKO model, and 38 DEGs for OPCs and 29 DEGs for microglia in the Ip3r2 KO model, 20-50% of the astrocyte DEG level. This suggests astrocytes are more sensitive to neuronal activity changes, as well as more profoundly affected by silencing their calcium activity. GO terms analysis of astrocyte DEGs in the VGlut2 cKO model revealed enrichment for pathways related to microtubule activity, axonal elongation and transport in the upregulated genes, and regulation of intracellular signal transduction related pathways in the downregulated genes (Fig S7A, Table S6D). In the Ip3r2 KO model, upregulated genes are enriched for signal transduction and response to alcohol, while downregulated genes are enriched in cAMP metabolism and pathways related to cellular biosynthesis processes (Fig S7B, Table S6D).

**Figure 7.**
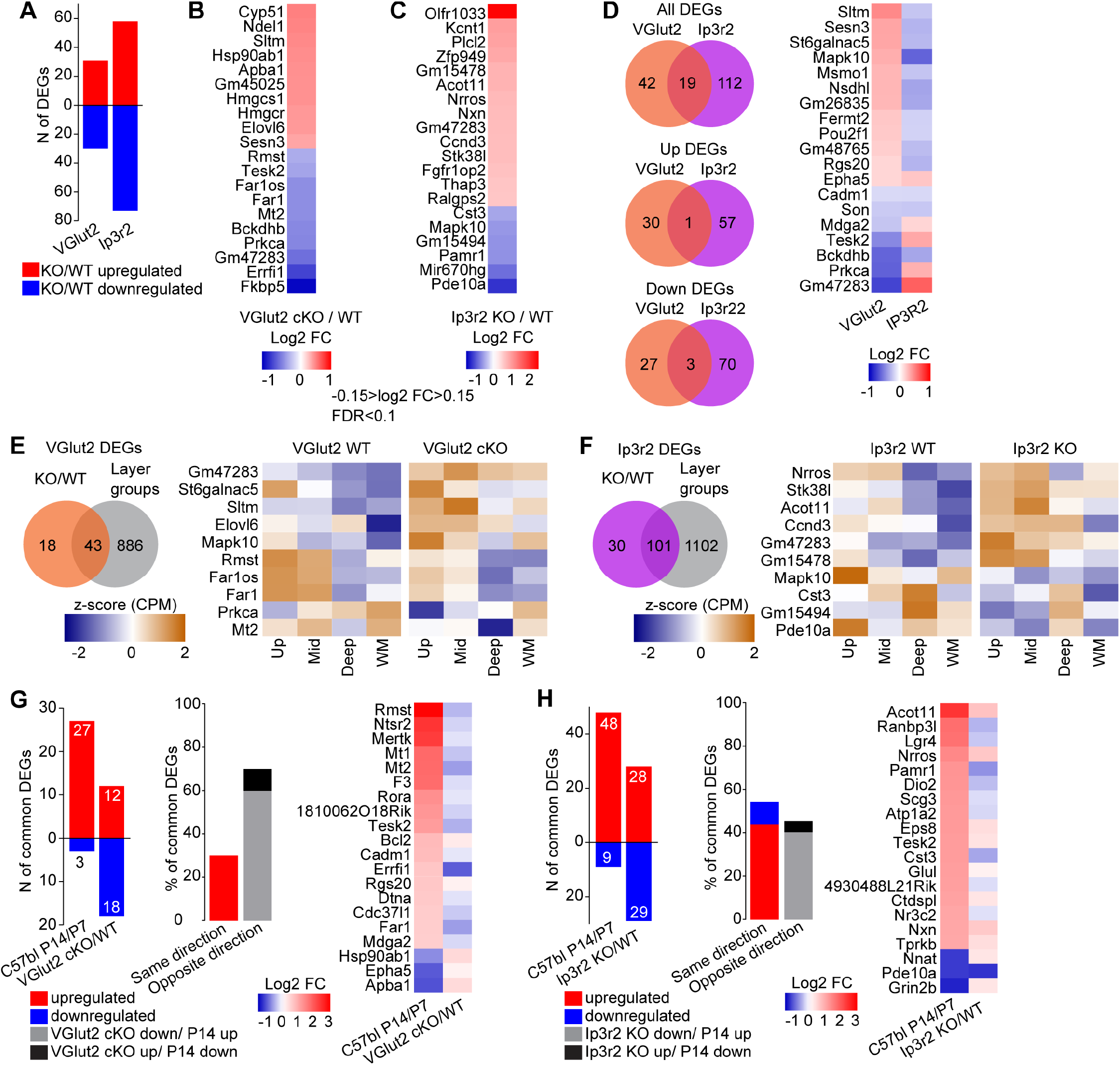
Global astrocyte gene expression changes following silencing of neuronal or astrocyte activity. See also Fig S7, Table S6. **A.** Number of DEGs identified for each model: VGlut2 cKO; 61 total DEGs, Ip3r2 KO; 131 total DEGs. Red – upregulated; Blue – downregulated. **B, C.** Heatmap showing top 20 DEGs identified in each model (B, VGlut2 cKO; C, Ip3r2 KO). Colors represent Log2 Fold change (FC) between each condition. Criteria for DEG selection: Log2 FC between −0.15 and 0.15; FDR <0.1. **D.** Venn diagrams showing DEGs common to both models. Heatmap shows FC of the 19 common DEGs. Most common DEGs are differentially regulated in each model (upregulated in VGlut2 cKO and downregulated in Ip3r2 KO). **E.** Venn diagram showing DEGs common to the VGlut2 cKO vs WT comparison and genes enriched in astrocyte layer groups. Heatmap of expression level z score of a select list of 10 genes, shows dysregulation of layer enrichment in the cKO mice compared to WT. Z-score was calculated for each gene using the combined data for WT and cKO average and standard deviation. **F.** Same analysis as E, but for the Ip3r2 KO model. **G.** Comparison between DEGs identified in the VGlut2 cKO dataset with DEGs between P7 and P14 of WT mice, identified in the bulk RNAseq dataset. A total of 30 VGlut2 cKO DEGs were also significantly up- or down-regulated at P14 vs P7. The majority of common DEGs were differentially regulated as shown in the bar graph and heatmap on the right. **H.** Same analysis as in G, but for the Ip3r2 KO model. A total of 57 DEGs were commonly identified in the developmental dataset. About half of the genes were commonly regulated, while the other half were differentially regulated as shown in the bar graph and heatmap on the right. For this analysis the selection criteria of Ribotag P7-P14 DEGs are FPKM>1; FDR<0.1.

As our smFISH analysis of Gpc4 and Chrdl1 identified that they are regulated in opposite directions by neuronal and astrocyte activity (Figs 4,5), we asked whether the DEGs identified by snRNAseq are similarly regulated. We found a total of 19 DEGs that are significantly altered in both the VGlut2 cKO and Ip3r2 KO models (Fig 7D). Only 1 gene was commonly upregulated, and only 3 genes commonly downregulated (21% of total common DEGs), while the remaining 15 genes showed opposing changes (79% of common DEGs; Fig 7D heatmap). The same effect of opposing changes was observed when the overlap between the enriched GO terms was compared (Fig S7C). For example, the term “intracellular signal transduction” is enriched in the downregulated gene list in the VGlut2 cKO model, whereas in the Ip3r2 KO model it is enriched in the upregulated gene list (Table S6D). Thus, opposing effects of neuronal and astrocyte activity extend beyond synapse-regulating genes.

Since we found that neuronal activity is important for regulating layer-specific expression of Gpc4, Gpc6 and Chrdl1 (Figs 3,4), we next asked if this is true for other DEGs identified in the snRNAseq dataset (Fig 7A-C). To address this we compared the list of neuronal or astrocyte activity-regulated DEGs with the list of layer enriched genes from the WT (Fig 6E,F, S6F, Table S5A-B). This identified 43 common genes in the VGlut2 cKO model (70% of all cKO/WT DEGs; Fig 7E), and 101 common DEGs in the Ip3r2 model (77% of all KO/WT DEGs; Fig 7F). Of these, some genes show a dysregulated layer-expression when activity is altered, similar to the findings described in Fig 4. For example, expression of Mapk10 is lowest in WM astrocytes in the WT, however in astrocytes from the VGlut2 cKO Mapk10 is upregulated in WM astrocytes compared to deep. Similarly, the expression of Pde10a is higher in the upper astrocyte group than other groups in WT, while in the Ip3r2 KO the level in that group is now low. Nevertheless, we also found many genes that while having different expression levels than WT after activity manipulation, still maintained their layer-enriched expression. For example, WT expression of the gene Gm47283 is higher in the upper than deep layer group. In the VGlut2 cKO while Gm47283 is upregulated in all groups, the relative expression between layers is maintained (Fig 7E). A similar pattern was observed in the Ip3r2 KO model for the gene Stk38I (Fig 7F). Thus, perturbation of neuronal or astrocyte activity influences the layer enrichment of some but not all genes, without causing gross rearrangement of overall astrocyte spatial identity.

Is neuronal and/or astrocyte activity necessary for regulating developmental changes in astrocytic gene expression? Our spatial smFISH analysis showed that developmental changes in Gpc4, Gpc6 and Chrdl1 are attenuated by blocking neuronal activity, resulting in opposite expression levels in VGlut2 cKO mice to those seen during normal development (Fig 4). On the other hand, astrocyte activity does not appear to regulate developmental astrocyte gene expression, with some genes in the Ip3r2 KO changing in the same direction as development (Gpc4, Chrdl1), with no effect on others (Gpc6) (Fig 5). To test if this is a general phenomenon beyond synapse-regulating genes we identified DEGs between P7 and P14 in our bulk RNAseq data (FC P14/P7 Fig 7G,H, Table S6C). We compared the DEG lists of VGlut2 cKO and Ip3r2 KO models against the genes that are significantly up- or down-regulated at P14 compared to P7 in the bulk data, in search for common genes (Fig 7G,H). This identified 30 DEGs from the VGlut2 cKO dataset (52% of all cKO/WT DEGs, Fig 7G), and 57 genes from the Ip3r2 KO dataset (44% of all KO/WT DEGs, Fig 7H). Comparing the direction of expression changes between normal development and the VGlut2 cKO DEGs, we observed that 70% of all common genes were regulated in the opposite direction (Fig 7G), similar to results found for Gpc4, Gpc6 and Chrdl1 (Fig 4), suggesting strong dependence of astrocyte developmental maturation on neuronal cues. On the other hand, Ip3r2 KO data showed 50% of genes displaying the same directionality as during normal development, and the other 50% showing the opposite regulation (Fig 7H). Therefore during development astrocyte-neuron communication enacts global gene expression changes in astrocytes, going beyond synapse-regulating genes.

## Discussion

In this study we demonstrate how astrocytes and neuronal synapses develop together in the postnatal brain, and how each cell signals to the other to ensure correct development. In particular, we show that: ● Astrocytes are unevenly distributed across the visual cortex, constituting more than 50% of all cells in L1, and about 10% of all cells in L2-6. ● Astrocyte transcriptome changes during development are correlated to expression changes in synaptic proteins. ● Astrocytes form heterogeneous populations in the developing cortex, based on their spatial location. ● Expression of select synapse-regulating genes (Gpc4, Gpc6 and Chrdl1) is differentially regulated during development at both temporal and spatial levels. ● Expression of astrocyte synapse-regulating genes is affected by changes in thalamic neuronal activity and astrocyte calcium activity. ● Neuronal and astrocyte activity regulate multiple non-overlapping genetic programs in astrocytes, demonstrating effects beyond synapse regulation.

### Astrocyte number and transcriptome alterations across development coincide with stages of synapse development

In the mouse cortex astrocytes begin to be generated right before birth and populate the cortex throughout the first month of life (Farhy-Tselnicker and Allen, 2018; Ge et al., 2012). During this time many changes are occurring in astrocytes, as well as in the synapses between neighboring neurons. We observed that the most significant change in astrocytes at the transcriptome level occurred between the first and second postnatal weeks (Fig 2). Similarly, an analysis of the synaptic proteome during development showed the largest difference between P9 and P15 (Gonzalez-Lozano et al., 2016), suggesting similar or overlapping regulatory mechanisms in both astrocytes and neurons. Our in-depth analysis of the developmental changes in the expression levels of the major components of glutamatergic synapses, presynaptic VGlut1, 2 and postsynaptic GluA1, 2 (Fig 1) revealed divergent expression profiles. While VGlut1 and GluA2 are strongly upregulated between the first and second postnatal weeks, VGlut2 and GluA1 exhibit a more gradual increase. This suggests different regulatory mechanisms at the level of individual pre- and post-synaptic proteins. Our observations fit with previous reports showing calcium-permeable AMPARs such as GluA1-containing are associated with immature synapses and expressed earlier than GluA2, which is inserted into the synapse at later ages marking a mature synapse (Brill and Huguenard, 2008).

In addition to the transcriptomic changes astrocyte numbers are also strongly regulated during development. Highly proliferative for the first two postnatal weeks (Ge et al., 2012), astrocytes rapidly expand and populate the entire cortex, peaking in numbers at P21 (Fig 1), showing a ~5-60 fold increase in numbers across the visual cortex at P21 compared to P1 (Table S1). Indeed, genes upregulated at P14 are uniquely enriched in GO terms related to cell proliferation and migration (Table S2B). Interestingly, the density of astrocytes remains fairly constant throughout development, suggesting their expansion rate is correlated with the overall expansion of the cortex. The mechanisms that regulate these migration patterns are still unknown, and seem to be largely unaffected by neuronal or astrocyte activity, as evident from the similar numbers of astrocytes within each cortical layer in both neuronal and astrocyte activity manipulation models tested here (Fig S4, S5). Future studies will determine the factors, or sets of factors that regulate the number and location of astrocytes within defined domains.

### Developmental expression of astrocyte genes is regulated by both neuronal and astrocyte activity

Ever since the astrocyte-derived factors that promote synapse formation were identified, an outstanding question in the field has been, how are they regulated (Baldwin and Eroglu, 2017; Farhy-Tselnicker and Allen, 2018)? Is it an astrocyte-intrinsic process, or is it affected/driven by changes in neuronal activity which occur as synapses develop? Our *in vitro* work together with previously published studies has provided evidence that neuronal activity can influence astrocyte gene expression and function at the synapse (Fig S4A, (Benediktsson et al., 2012; Bernardinelli et al., 2014b; Durkee and Araque, 2019; Hasel et al., 2017)). However, whether neuronal activity regulates astrocytes in the developing brain *in vivo* has not been systematically addressed. Here, silencing thalamic neurons that project to the VC by knocking out VGlut2 resulted in attenuation of the developmental expression changes in astrocyte genes, as well as AMPAR subunits, at P14 but not at P7. This suggests a delay in circuit maturation, similar to other studies employing visual deprivation methods, and observing delayed maturation of VC neurons (Albanese et al., 1983; Desai et al., 2002; Freire, 1978; Funahashi et al., 2013; Ishikawa et al., 2014; Ko et al., 2014), and astrocyte morphology (Müller, 1990; Stogsdill et al., 2017). Furthermore, this regulation goes beyond synapse development, as snRNaseq analysis identified many genes differentially expressed by astrocytes in VGlut2 cKO mice which are related to additional cellular processes. Notably, VGlut2 cKO did not affect the levels of VGlut1 (Fig 4; (Wallén-Mackenzie et al., 2010)), which marks cortico-cortical connections, suggesting the normal upregulation in VGlut1 that occurs at P14 is either intrinsic to the cortical neurons, and/or regulated by other mechanisms than dLGN-VC evoked activity.

Our study revealed an additional important layer of regulation of astrocyte expression of Gpc4 and Chrdl1, and that is by Ip3r2-mediated astrocyte calcium activity (Petravicz et al., 2014; Srinivasan et al., 2015). Interestingly, silencing the ability of astrocytes to increase intracellular calcium by knocking out Ip3r2 resulted in an opposite regulation of gene expression to the ones observed in the VGlut2 cKO mice, and did not correspond to layer-specific developmental changes, suggesting a more global role of astrocyte calcium activity in the regulation of Gpc4 and Chrdl1 gene expression. Notably, Gpc6 expression was unaltered in Ip3r2 KO VC at P14, suggesting distinct regulation of expression of the two glypican family members. snRNAseq analysis identified many additional genes regulated by astrocyte calcium activity, both synapse and non-synapse related, however these weren’t correlated with developmental changes. These results suggest that astrocyte Ca^2+^ activity is an important intrinsic mechanism for regulating developmental gene expression, but is not tied to changes occurring following eye opening.

### Astrocytes form diverse populations in the developing mouse visual cortex

The diversity of neurons based on location, morphology, connectivity and activity patterns has been extensively studied for decades, with multiple subtypes of excitatory and inhibitory neurons identified (Kepecs and Fishell, 2014; Migliore and Shepherd, 2005; Zeisel et al., 2015). For a long time cortical protoplasmic astrocytes were viewed as a homogeneous population. However recent studies investigating astrocyte heterogeneity using both bulk and single-cell sequencing have shown that within the cortex, astrocytes are also heterogeneous (Batiuk et al., 2020; Bayraktar et al., 2020; Lanjakornsiripan et al., 2018). Unlike neurons, astrocytes do not fall into the 6-layer categories, but rather exist on a gradient of transcriptomically separable yet overlapping groups. Indeed, our snRNAseq data shows that the biggest differences are between astrocytes of the upper and deep layer groups (L1-2/3 vs L5-6), while upper and mid-layer groups (L1-2/3-L2/3-5) are the most similar. Nevertheless, we have identified several astrocyte population marker genes (such as Dcc, Siah3 or Kcnd2; Fig 6), that are significantly enriched in one group over others. In the future, these could be used to target specific populations of astrocytes, similar to the methods employed for neurons, in order to manipulate astrocytes that interact with specific synapse types or circuits. Importantly, while blocking thalamo-cortical activity did alter the expression of numerous genes in astrocytes, it did not alter the layer patterning of the cells, showing that this is not a major factor in driving layer-enriched gene expression. Indeed altering the identity of local cortical neurons by using Dab1 KO mice, in which cortical layer neurons are reversed, does alter astrocyte layer identity, suggesting a role for local cues (Lanjakornsiripan et al., 2018). Our findings further suggest that neuronal activity acts to fine-tune the level of astrocyte genes that are important for neuronal function, rather than determining their presence or absence.

Interestingly some of the synapse-regulating genes we profiled display layer-specific expression changes across development (Fig 3). We found a correlation between Chrdl1 upregulation in the upper layers with that of GluA2, consistent with our previous findings regarding Chrdl1 regulation of GluA2 levels (Blanco-Suarez et al., 2018). A more complex picture emerges for Gpc4, Gpc6, and GluA1, the AMPAR subunit regulated by these genes (Allen et al., 2012), as they do not show such correlation in their expression patterns. GluA1 protein levels steadily increase across development, peaking in most layers at P7, and do not show downregulation at P14 in L1 (as was observed for Gpc4), or upregulation in deeper layers (as was shown for Gpc6). Still, changes in Gpc4 expression are contributing to the levels of GluA1, as GluA1 is affected in correlation with changes in Gpc4 expression in the neuronal and astrocyte activity deficit models (Figs 4,5), and GluA1 levels are reduced in the VC of Gpc4 KO mice (Farhy-Tselnicker et al., 2017). One possibility is that Gpc4 and Gpc6 may regulate GluA1 levels at specific synapses, such as glutamatergic terminals onto interneurons in L1, or deep layer cortical neurons, making it hard to distinguish their specific effect when analyzing synapses as a group.

This study demonstrates that the correct formation of synapses and hence neuronal circuit connectivity depends on precise communication between neurons and astrocytes, where disruption in one cell type leads to disruption in the other and an overall dysregulation of synapse formation. It further shows that astrocyte regulation of synapses is intimately linked to environmental changes. Thus, an image of astrocyte identity emerges as highly plastic and dynamic cells, actively perceiving and responding to their environment. Future studies to determine the precise nature of astrocyte plasticity and to further distinguish intrinsic and extrinsic influences on these cells will give insight into their function in both health and disease.

## ACKNOWLEDGEMENTS

We thank members of the Allen lab for helpful discussions on the project. This work was supported by NIH NINDS grants to NJA: NS105742 and NS089791. Work in the lab of N.J.A. is supported by the Hearst Foundation, the Pew Foundation, and the CZI Neurodegeneration Network. This work was supported by Core Facilities of the Salk Institute (Next Generation Sequencing, Bioinformatics, Biophotonics: NIH NCI CCSG P30 014195, the Waitt, Helmsley and Chapman Foundations). J.R.E is an Investigator of the Howard Hughes Medical Institute.

## AUTHOR CONTRIBUTIONS

I.F.-T. and N.J.A conceived the project, designed experiments, analyzed data and wrote the manuscript, with input from other authors. I.F.-T., M.M.B., C.D., E.B-S. performed experiments and analyzed data. C.F., H.L., G.A.E. analyzed data, with supervision from M.S. and J.R.E.

## SUPPLEMENTAL INFORMATION

Supplemental information includes seven figures and six tables.

## Supplemental Table Legends

**Table S1 (related to Figures 1, S1). Development of astrocytes and synapses in the mouse visual cortex.** Full statistical analysis of astrocyte numbers, VGlut1, VGlut2, GluA1, and GluA2 changes during development. Each comparison (e.g. astrocyte number/area, comparison between ages within each layer) are labeled accordingly. Statistical comparison between ages within each layer (top), as well as between layers at each age (bottom) are shown.

**Table S2 (related to Figures 2, S2). Determination of the astrocyte transcriptome across visual cortex development. A.** Complete list of genes (expression levels shown as FPKM) at each developmental stage as indicated. Expression levels for each sample, as well as average are shown, as well as pairwise analysis and FDR values. **B.** Complete list of GO terms (Biological Process) identified for astrocyte enriched genes at each developmental time point as indicated.

**Table S3 (related to Figures 3, S3). Synapse-regulating genes in astrocytes show differential spatio-temporal expression. A.** Full statistical analysis of developmental changes in mRNA expression of selected synapse regulating genes by smFISH. Averages and analysis calculated for N=3, i.e. per mouse. **B.** Full statistical analysis of developmental changes in mRNA expression of selected synapse regulating genes by smFISH. Averages and analysis calculated for n=~50-350, i.e. total number of astrocytes per group (across the 3 mice). Statistical comparison between ages within each layer (top), as well as between layers at each age (bottom) are shown.

**Table S4 (related to Figures 4, S4, 5, S5). Neuronal and astrocyte activity regulate astrocyte expression of synapse regulating genes. A.** Full statistical analysis of mRNA expression differences between WT and KO in VGlut2 and Ip3r2 models. Averages and analysis calculated for N=5, i.e. per mouse. **B.** Full statistical analysis of mRNA expression differences between WT and KO in VGlut2 and Ip3r2 models. Averages and analysis calculated for n=~200-400, i.e. total number of astrocytes per group (across 5 mice). All comparisons are between WT and KO within each layer.

**Table S5 (related to Figures 6, S6). Unbiased determination of astrocyte layer-enriched genes. A.** Complete list of DEGs identified in pairwise analysis between astrocyte layer groups for VGlut2 WT dataset. **B.** Complete list of DEGs identified in pairwise analysis between astrocyte layer groups for Ip3r2 WT dataset. **C.** Complete list of GO terms (Biological Process) identified for astrocyte layer group enriched genes for the VGlut2 WT dataset.

**Table S6 (related to Figures 7, S7). Global astrocyte gene expression changes following silencing of neuronal or astrocyte activity. A.** Complete list of DEGs between WT and KO for each model, VGlut2 and Ip3r2. **B.** Complete list of genes common to KO/WT DEGs and layer group enriched DEGs identified for the WT. **C.** Complete list of genes common to KO/WT DEGs and developmentally regulated genes (P14/P7 DEGs) identified in bulk RNAseq. **D.** Complete list of GO terms (Biological Process) identified for DEGs between WT and KO (VGlut2, Ip3r2), as well as terms identified for DEGs common to both models.

## METHODS AND MATERIALS

### ANIMALS

All animal work was approved by the Salk Institute Institutional Animal Care and Use Committee.

#### Rats

Sprague Dawley rats (Charles Rivers) were maintained in the Salk Institute animal facility, under a 12 hour light:dark cycle with ad libitum access to food and water. Rat pups (both male and female) were used at postnatal day (P) 1-2 for preparation of primary cortical astrocyte cultures, and at P5-P7 for preparation of purified immunopanned retinal ganglion cell (RGC) neuronal cultures.

#### Mice

Mice were maintained in the Salk Institute animal facility, under a 12 hour light:dark cycle with ad libitum access to food and water. Both male and female mice were used for experiments. The following mouse lines were used:

***Wild-type (WT; C57Bl6/J)*** were purchased from Jackson Labs and bred in-house (Jax #000664). Mice were used for breeding and backcrossing, and as non-littermate controls.
***Ribotag floxed* (B6N.129-Rpl22tm1.1Psam/J)** were obtained from Jackson Labs (Jax #011029). Mice were maintained as homozygous for floxed Rpl22 on C57Bl6/J background, and crossed to mice expressing cre recombinase for experiments.
***Gfap-cre* (B6.Cg-Tg (Gfap-cre)73.12Mvs/J)** mice were obtained from Jackson Labs (Jax #012886), and bred in house to generate cre+ females.
*To generate Astrocyte-ribotag mice* homozygous flox-Rpl22-HA males were crossed to Gfap-cre hemizygous females. Male mice hemizygous for cre and heterozygous for flox-Rpl22-HA (Rpl22-HA+; Gfapcre+) were used for all experiments.
***Aldh1l1-GFP* (Tg(Aldh1l1-EGFP)OFC789Gsat/Mmucd)** were obtained from MMRRC. They were backcrossed to C57Bl6/J background (Jax: 000664) for at least 4 generations prior to conducting experiments.
***VGlut2 floxed* (Slc17a6tm1Lowl/J)** were obtained from Jackson Labs (Jax #012898). Mice were maintained as homozygous for floxed VGlut2 on a C57Bl6/J background, and crossed to mice expressing cre recombinase for experiments.
***RORα-IRES-Cre*** were obtained from Dennis O’Leary at the Salk Institute, and described in (Chou et al., 2013) and (Farhy-Tselnicker et al., 2017). Mice were backcrossed to C57Bl6/J (Jax #000664) for 3-4 generations before being crossed to the VGlut2 flox line. Expression of cre recombinase in the thalamus was confirmed by crossing the RORα cre mouse with a tdTomato reporter mouse line (Jax #007914) (Fig 4A).
*To generate conditional VGlut2 KO mice* for experiments, VGlut2^f/f^;cre-ve females were bred to VGlut2^f/f^;RORα cre positive males. Homozygous VGlut2 flox;RORα cre positive littermates were compared with homozygous VGlut2 flox;RORα cre −ve in each experiment.
*To generate het and homozygous VGlut2 cKO mice expressing a fluorescent reporter in the recombined neurons* (Fig 4, S4), VGlut2^f/+^;RORα cre positive; tdTomato positive males were crossed to VGlut2^f/f^ females. As control in these experiments, RORα cre positive mice were crossed to tdTomato positive, to generate VGlut2^+/+^RORα;cre positive; tdTomato positive mice. ***IP3R2 KO*** was obtained from Ju Chen lab at UCSD (Li et al., 2005) and maintained on C57BL6/J background, either as KO x KO breeding scheme, or het x het breeding scheme. Both littermate and non-littermate pairs of WT and KO mice were used for experiments. For non-littermate pairs, C57Bl6/J that were bred in-house were used as control.
In all cases, when littermates could not be used as control, mice were matched by age, size, fur color and condition, and eye opening to ensure identical developmental stage.

#### Mouse Tissue collection

Tissue was collected at the following developmental time points: post-natal day (P) 1, P4, P7, P14, P21, P28, and P120.

##### Ribotag RNAseq

All mice were collected between 9:30am and 12:30pm on the day of experiment. Mice were anesthetized by I.P. injection of 100 mg/kg Ketamine (Victor Medical Company)/20 mg/kg Xylazine (Anased) mix, and transcardially perfused with 10ml PBS then 10ml 1% PFA. Brains were dissected in 2.5mM HEPES-KOH pH 7.4, 35mM glucose, 4mM NaHCO3 in 1x Hank’s Balanced Salt Solution with 100μg/ml cycloheximide added fresh (Heiman et al., 2014). Brains were cut at approximately bregma −2.4 to isolate the visual cortex, the cortex was carefully detached from the subcortical areas, and any visible white matter was removed. Lateral cuts were made at 1mm and 3mm from the midline to further isolate the VC section, and Ribotag pulldown was immediately performed. For each time point the visual cortices from 2 mice (Rpl22-HA+; Gfap cre+) were pooled for RNA isolation and RNA sequencing library preparation. P7 = 3 biological replicates (6 mice, 2 × 3); P14 = 4 biological replicates (8 mice, 2 × 4); P28 = 5 biological replicates (10 mice, 2 × 5); P120 = 6 biological replicates, 3 new samples (6 mice, 2 × 3), plus for data analysis 3 additional P120 biological replicates from a previously published study from the lab (Boisvert et al 2018; GEO GSE99791), collected and processed in the same way, were included to increase the power of the analysis.

##### Histology (smFISH In situ hybridization and Immunostaining)

Mice aged P4 and older were anaesthetized by I.P. injection of 100 mg/kg Ketamine (Victor Medical Company)/20 mg/kg Xylazine (Anased) mix and transcardially perfused with PBS, then 4% PFA at room temp. Brains were removed and incubated in 4% PFA overnight at 4C, then washed 3X 5 min with PBS, and cryoprotected in 30% sucrose for 2-3 days, before being embedded in TFM media (General data healthcare #TFM-5), frozen in dry ice-ethanol slurry solution, and stored at −80C until use. P1 mice were decapitated and brains removed without perfusion, briefly washed in PBS and put in 4% PFA overnight at 4C, followed by a similar procedure as described above for older mice. Brains were sectioned using a cryostat (Hacker Industries #OTF5000) in sagittal or coronal orientations depending on experimental needs at a slice thickness of 16-25 μm. Sections were mounted on Superfrost plus slides (Fisher #1255015). Immunostaining for synaptic markers and smFISH were performed on the same day of sectioning. 3-5 mice were used for each experimental group. For each mouse, 3 sections were imaged and analyzed.

##### Single nucleus RNAseq and Western blot

Mice were anesthetized by I.P. injection of 100 mg/kg Ketamine (Victor Medical Company)/20 mg/kg Xylazine (Anased) mix, then decapitated. Brains were rapidly removed and the visual cortex dissected in ice-cold PBS using the same coordinates as described for Ribotag RNAseq. Dissected cortices were snap frozen, and kept at −80C until use. For snRNAseq, 4 mice were collected for each experimental group. For Western blot, 2-4 independent experiments/samples for each condition were analyzed.

### RNAseq

#### Bulk RNAseq using Ribotag

##### Ribotag pulldown

A modified Ribotag protocol was performed to isolate astrocyte enriched RNA. Briefly, brain samples were homogenized using a Dounce homogenizer (Sigma #D9063) in 2ml cycloheximide-supplemented homogenization buffer (1% NP-40, 0.1M KCl, 0.05M Tris, pH 7.4, 0.012M MgCl2 in RNase free water, with 1:1000 1M DTT, 1mg/mL heparin, 0.1mg/mL cycloheximide, 1:100 Protease inhibitors, and 1:200 RNAsin added fresh). Homogenates were centrifuged and the supernatant incubated on a rotator at 4°C for 4 hours with 5ul anti-HA antibody to bind the HA-tagged ribosomes (CST Rb anti-HA #3724, 1:200). Magnetic IgG beads (Thermo Scientific Pierce #88847) were conjugated to the antibody-ribosome complex via overnight incubation on a rotator at 4°C. Samples were washed with a high salt buffer (0.333M KCl, 1% NP40, 1:2000 1M DTT, 0.1mg/mL cycloheximide, 0.05M Tris pH 7.4, 0.012M MgCl2 in RNase-free water), and RNA released from ribosomes with 350uL RLT buffer (from Qiagen RNeasy kit) with 1% BME. RNA was purified using RNeasy Plus Micro kit (Qiagen 74034) according to manufacturer instructions and eluted into 16ul RNase-free water. Eluted RNA was stored at −80°C. For each time point, 50ul of homogenate (pre- anti-HA antibody addition) was set aside after centrifugation, kept at −20°C overnight, and purified via RNeasy Micro kit as an ‘input’ sample, and used to determine astrocyte enrichment.

##### Library generation and sequencing

RNA quantity and quality were measured with a Tape Station (Agilent) and Qubit Fluorimeter (ThermoFisher) before library preparation. >100ng of RNA was used to make libraries. mRNA was extracted with oligo-dT beads, capturing polyA tails, and cDNA libraries made with Illumina TruSeq Stranded mRNA Library Preparation Kit (RS-122-2101) by the Salk Institute Next Generation Sequencing (NGS) Core. Samples were sequenced on an Illumina HiSeq 2500 with single-end 50 base-pair reads, at 12-70 million reads per sample.

##### RNA sequencing mapping, analysis, and statistics

Raw sequencing data was demultiplexed and converted into FASTQ files using CASAVA (v1.8.2), and quality tested with FASTQC v0.11.2. Alignment to the mm10 genome was performed using the STAR aligner version 2.5.1b (Dobin et al., 2013). Mapping was carried out using default parameters (up to 10 mismatches per read, and up to 9 multi-mapping locations per read), and a high ratio of uniquely mapped reads (>75%) was confirmed with exonic alignment inspected to ensure that reads were mapped predominantly to annotated exons. Raw and normalized (FPKM) gene expression was quantified across all genes (RNAseq) using the top-expressed isoform as a proxy for gene expression using HOMER v4.10 (Heinz et al., 2010), resulting in 10-55 million uniquely mapped reads in exons. Principal Component Analysis was carried out with prcomp in R 3.4.3 on normalized counts. Differential gene expression was carried out using the DESeq2 (Love et al., 2014) package version 1.14.1 using the HOMER getDiffExpression.pl script with default normalization and using replicates to compute within-group dispersion. Significance for differential expression was defined as adjusted P<0.05, calculated using Benjamini-Hochberg’s procedure for multiple comparison adjustment. Significantly altered genes are presented in 3 categories:

*All genes:* FPKM >1, adjusted p<0.05
*Astrocyte-expressed genes:* ribotag pulldown (astrocyte)/input (all cells) >0.75, FPKM >1, adjusted p<0.05
*Astrocyte-enriched genes:* ribotag pulldown (astrocyte)/input (all cells) >3, FPKM >1, adjusted p<0.05
See also (Boisvert et al., 2018).

##### GO enrichment analysis

GO terms that are enriched in astrocytes at each developmental stage, were identified using the String database (https://string-db.org/)(Szklarczyk et al., 2019). A search using multiple proteins by gene name was performed with the default parameters, and GO Biological Process category selected and exported from the analysis tab. GO terms with gene ratio above 0.5 were selected, and plotted for each age group using dot charts (function dotchart in R), with x-axis showing the ratio of genes overlapping with each GO term, and dot size is the significance of the overlap (adj. P value). GO terms common to all age groups were obtained using the Venn Diagram (http://www.interactivenn.net/) (Heberle et al., 2015), and terms with gene ratio above 0.5 were selected, and plotted using dot charts. A full list of GO terms is presented in Table S2B.

#### Single nucleus RNAseq

##### Sample preparation

A total of 8 samples (2 for VGlut2 WT, 2 for VGlut2 cKO, 2 for Ip3r2 WT, 2 for Ip3r2 KO) were sequenced to obtain the dataset described in Figs 6-7. The samples were as follows: VGlut2 WT_1; VGlut2 cKO_1; VGlut2 WT_2; VGlut2 cKO_2; Ip3r2 WT_1; Ip3r2 KO_1; Ip3r2 WT_2; Ip3r2 KO_2. Each group consisted of 1 replicate from male mice and 1 replicate from female mice. Each replicate consisted of the VC from both hemispheres of 2 mice of the same genotype and gender. Nuclear isolation, FACS sorting, 10x Barcoding and cDNA preparation were performed on the same day using 1 WT and KO pair, that were processed in parallel, resulting in 4 separate procedures. cDNA was stored at −20C until all samples were collected. Library preparation and sequencing were carried out at the same time for all 8 samples.

##### Nuclei preparation

Nuclei were isolated from frozen visual cortex tissue. Tissue was manually homogenized using a 2 step Dounce homogenizer (A and B) (Sigma #D9063) in NIMT buffer, containing (in mM: 250 Sucrose, 25 KCl, 5 MgCl2, 10 Tris-Cl pH 8, 1 DTT; 1:100 dilution of: Triton X100, Protease Inhibitor Cocktail (Sigma #P8340); and 1:1000 dilution of: RNaseOUT™ Recombinant Ribonuclease Inhibitor (Thermo #10777019); SUPERase• In™ RNase Inhibitor (Thermo #AM2694)) on ice. Homogenized samples were mixed with 50% Iodixanol (OptiPrep™ Density Gradient Medium (Sigma #D1556)) and loaded onto 25% Iodixanol cushion, and centrifuged at 10,000g for 20 min at 4C in a swinging bucket rotor (Sorval HS-4). Pellets resuspended in ice-cold DPBS (HyClone) with 1:1000 dilution of: RNaseOUT™ Recombinant Ribonuclease Inhibitor (Thermo #10777019); SUPERase• In™ RNase Inhibitor (Thermo #AM2694). Nuclei were then incubated for 7 min on ice with Hoechst 33342 Solution (20 mM) (Thermo #62249) (final concentration 0.5μM), followed by centrifugation at 1000g for 10min at 4C to pellet nuclei. Pellets were resuspended in blocking buffer containing DPBS with RNAse inhibitors, and 1:10 dilution of pure BSA, and blocked for 30min on ice. Neun-Alexa488 pre-conjugated antibody (Millipore #MAB377X) was then added at 1:1000 dilution, and incubated for at least 1 hr on ice before proceeding to Flow cytometry sorting.

##### Flow cytometry

Fluorescence-Activated Nuclei Sorting (FANS) was performed in the Salk Institute Flow Cytometry core using a BD FACS Aria Fusion sorter with PBS for sheath fluid (a 100-μm nozzle was used for these experiments with sheath pressure set to 20 PSI). Hoechst-positive nuclei were gated first (fluorescence measured in the BV421 channel), followed by exclusion of debris using forward and side scatter pulse area parameters (FSC-A and SSC-A), exclusion of aggregates using pulse width (FSC-W and SSC-W), before gating populations based on NEUN fluorescence (using the FITC channel). To isolate the non-neuronal cell population, nuclei devoid of FITC signal (Neun-) were collected (Fig 6A). Nuclei were purified using a 1-drop single-cell sort mode (for counting accuracy); these were directly deposited into a 1.5 ml eppendorf without additional buffer (to yield a sufficient concentration that permitted direct loading onto the 10x chip).

Sorted NeuN-negative nuclei were immediately processed with 10x Chromium kit (10x Genomics) for single nucleus barcoding. Nuclei were kept on ice for the entire process. At each time, WT and KO samples were processed in parallel on the same day.

##### 10x Chromium barcoding, library preparation and sequencing

Single nuclei separation, barcoding, and cDNA generation were performed following the manufacturer’s instruction using the Chromium single cell 3’ kit (V3, 10x genomics PN-1000073). cDNA concentration and quality were measured using Qubit Fluorimeter (ThermoFisher) and Tape Station (Agilent) respectively, and was stored at −20C until library preparation.

Libraries were generated from all samples at the same time (8 total samples, 2 WT/2 KO Vglut2cKO model; 2 WT/2KO IP3R2 KO model) following manufacturer’s instructions using the Chromium single cell 3’ kit (V3, 10x genomics PN-1000075). Library quality was assessed with a Tape station (Agilent). NovaSeq sequencing was performed at the UCSF Center for Advanced Technology, at ~300 million reads/ sample (60,000 reads/cell).

##### Single-cell RNA-seq Data Preprocessing and Clustering

Data was demultiplexed and mapped onto the mouse genome (mm10) using 10X Cellranger (v3.1.0) with default parameters. Cell barcodes with < 200 genes detected were discarded due to low coverage. Doublets were identified and removed using Scrublet (Wolock et al., 2019) with its default setting in each sample. The average number of UMIs per cell was 2310 +/− 878; average number of genes detected per cell (UMI >= 1) was 1168 +/− 328. Cell clusters were identified using Scanpy (v1.4.3), following the clustering process described in (Luecken and Theis, 2019). All the samples were combined and used the top 5000 highly variable genes as the input dimension reduction. To identify clusters, Scanorama (v1.0.0, default parameter, k=20) (Hie et al., 2019) was used to perform batch correction and dimension reduction (30 PCs), followed by Leiden clustering (Traag et al., 2019) (resolution = 1). Data was visualized using the UMAP embedding (McInnes et al., 2018) function from Scanpy. The ensemble clustering identified all astrocytes as one cluster, to further identify astrocytes subtypes, we repeated the same clustering process on the astrocytes cluster only and got four subtypes. Astrocyte clusters were annotated using cell-type marker genes identified from previous studies to label distinct cortical astrocyte populations (Bayraktar et al., 2020; Lanjakornsiripan et al., 2018; Marques et al., 2016; Tasic et al., 2018; Van Hove et al.; Zeisel et al., 2018).

##### Identifying Differential Expressed Genes (DEG)

To identify cluster-specific DEGs, we used the scanpy.tl.rank_gene_groups function to perform the Wilcoxon rank-sum test with Benjamini-Hochberg correction to compare cells from each cluster with the remaining cells. Genes with FDR < 0.1 and log2 fold change between −0.15 and 0.15 were identified as DEGs. To identify DEGs between KO and WT, we performed the same analysis using combined astrocyte clusters. All comparisons were performed separately for VGlut2 cKO, and Ip3r2 KO samples.

##### GO enrichment analysis

GO terms that are enriched in astrocyte gene groups within each cluster, as well as genes regulated by neuronal or astrocyte activity, were identified using the String database (https://string-db.org/)(Szklarczyk et al., 2019). A search using multiple proteins by gene name was performed separately on VGlut2 cKO and Ip3r2 KO samples, and up- and down-regulated DEGs, using the default parameters, and GO Biological Process category selected and exported from the analysis tab. A maximum of 25 GO terms (with lowest adj. p-value) were selected, and plotted for each model using dot charts (function dotchart in R), with x-axis showing the ratio of genes overlapping with each GO term, and dot size is significance of the overlap (adj. p-value). GO terms common to both models were obtained using the Venn Diagram (http://www.interactivenn.net/) (Heberle et al., 2015), and plotted using dot charts. A full list of GO terms is presented in Tables S5C, S6D.

### CELL CULTURE

#### Retinal Ganglion Cell (RGC) neuron purification and culture

RGC purification and culture was performed as described (Allen et al., 2012; Ullian et al., 2001; Winzeler and Wang, 2013). Briefly, retinas from P5-P7 rat pups of both sexes were removed and placed in DPBS (HyClone #SH30264). Retinas were digested with Papain (Worthington #PAP2 3176; 50 units) for 30 min at 34C, triturated with Low OVO (15 mg/ml trypsin inhibitor (Worthington #LS003086)), then High OVO (30 mg/ml trypsin inhibitor (Worthington #LS003086)) solutions. The cell suspension was then added to lectin (Vector #L-1100) coated Petri dishes to pull down microglia and fibroblast cells for 5-10 min at room temp. The remaining cells were then added to T11D7 hybridoma supernatant coated petri dishes for 40 min at room temp, which specifically binds RGCs. After washing off the non-binding cells with DPBS, pure RGCs were released by trypsin treatment (Sigma #T9935) to cleave cell-antibody bond, and collected. RGCs were plated on 6-well plates coated with PDL (Sigma # P6407) and laminin (Cultrex Trevigen #3400-010-01) at a density of 125,000 cells/well. RGCs were maintained in the following media: 50% DMEM (Life tech #11960044); 50% Neurobasal (Life Tech #21103049); Penicillin-Streptomycin (LifeTech #15140-122); glutamax (Life Tech #35050-061); sodium pyruvate (Life Tech #11360-070); N-acetyl-L-cysteine (Sigma #A8199); insulin (Sigma #I1882); triiodo-thyronine (Sigma #T6397); SATO (containing: transferrin (Sigma #T-1147); BSA (Sigma #A-4161); progesterone (Sigma #P6149); putrescine (Sigma #P5780); sodium selenite (Sigma #S9133)); and B27 (see (Winzeler and Wang, 2013) for recipe). For complete growth media, the media was supplemented with BDNF (Peprotech #450-02), CNTF (Peprotech #450-13), and forskolin (Sigma #F6886). The next day, half of the media was replaced with media containing FUDR (13 μg/μl final concentration Sigma #F0503) to inhibit fibroblast growth. Cells were fed by replacing half of the media with fresh equilibrated media every 3-4 days. RGCs were maintained at 37C/10%CO2 and kept in culture for at least 7 days prior to treatment to allow for full process outgrowth.

#### Astrocyte preparation and culture

Primary astrocytes from rat cortex were prepared as described (Allen et al., 2012; McCarthy and de Vellis, 1980). Briefly, the cerebral cortex from P1-P2 rat pups were removed and placed in DPBS (HyClone #SH30264). The meninges and hippocampi were removed and discarded. The remaining cortices were diced and digested with Papain (Worthington #LS003126; 330 units) for 1 hr and 15 min in 37C 10% CO2 cell culture incubator. Cells were triturated in Low OVO and then High OVO containing solutions, and plated in PDL coated 75cm tissue culture flasks. 3 days after plating, flasks were manually shaken to remove upper cell layers which contained mostly non-astrocytic cells. 2 days after shake off, ARA-C (10 μM final concentration; Sigma #C1768) was added for 48 hours to inhibit the other proliferating cells, which divide faster than astrocytes. Finally, astrocytes were plated in 15 cm cell culture plates coated with PDL at 2-3 million cells/dish and passaged once a week. Astrocytes were maintained at 37C/10%CO2 and kept in culture for 3-4 weeks. Astrocyte culture medium was DMEM (Life tech #11960044) supplemented with 10% Heat inactivated FBS (LifeTech #10437028), Penicillin-Streptomycin (LifeTech #15140-122), glutamax (Life Tech #35050-061), sodium pyruvate (Life Tech #11360-070), hydrocortisone (Sigma #H0888), and N-acetyl-L-cysteine (Sigma #A8199).

#### Treatment of astrocyte cultures with cultured neurons

Cultured astrocytes were plated on cell culture inserts (Falcon #353102) at 250,000 cells/insert. Inserts were added to 6 well plates containing either plated RGC neurons (at ~125,000 cells/ well), or empty wells coated with PDL and laminin (similar to RGC plated wells), and containing media. Cells were incubated together for 4 days in low protein conditioning media containing (50% DMEM, 50% Neurobasal media; Penicillin-Streptomycin; Glutamax and sodium pyruvate, NAC, BDNF, CNTF, forskolin), after which conditioned media was collected and concentrated 50-fold using 10 kDa cutoff concentrators (Sartorius #14558502). Protein concentration was measured using the Bradford assay. 3 experimental groups were compared: RGCs alone, astrocytes alone, astrocytes + RGCs.

### WESTERN BLOT

Samples were heated in reducing loading dye (Thermo # 39000) for 45 min at 55C. For conditioned media, 10 μg/ well was loaded; for tissue lysates 20 μg/ lane was loaded. Samples were resolved on 4-12% bis-tris or bolt gels (Invitrogen #NW04120) for 30-40 min at 150-200V. Proteins were transferred to PVDF membranes at 100V for 1 hr, then blocked in 1% casein (Biorad #1610782) in TBS (Bioworld #105300272) blocking buffer for 1 hr at room temp on a shaker. Primary antibodies were applied overnight at 4C diluted in blocking buffer. The antibodies used were: Rb anti-Glypican 4 (Proteintech #13048-1-AP; 1:500), Rb anti IP3R2 (a gift from Ju Chen lab, UCSD 1:1000), Ms anti-Tubulin (Thermo #MA5-16308 1:5000). The next day, membranes were washed 3X 10 min with TBS-0.1%Tween and the appropriate secondary antibody conjugated to Alexa fluor 680 was applied for 2 hrs at room temp (dilution 1:10,000). Bands were visualized using the Odyssey Infrared Imager (LiCor) and band intensity analyzed using the Image Studio software (LiCor).

### HISTOLOGY

#### Immunostaining in mouse brain tissue

The slides containing the sections were blocked for 1 hr at room temp in blocking buffer containing antibody buffer (100 mM L-lysine and 0.3% Triton X-100 in PBS) supplemented with 10% heat-inactivated normal goat serum. Primary antibodies diluted in antibody buffer with 5% goat serum were incubated overnight at 4C. The next day slides were washed 3X 5 min with PBS with 0.2% Triton X-100 and secondary antibodies conjugated to Alexa fluor were applied for 2 hrs at room temp. Slides were mounted with the SlowFade Gold with DAPI mounting media (Life Tech #S36939), covered with 1.5 glass coverslip (Fisher #12544E) and sealed with clear nail polish. The following antibodies were used: Chk anti GFP (Millipore #06-896, 1:500), Rb anti Sox9 (Abcam #ab185966, 1:2000), Rb anti Aldh1l1 (Abcam #ab-87117, 1:500), Rb anti HA (CST #3724), Rb anti S100β (Abcam #ab52642, 1:100), Ms anti Neun (Millipore #MAB377), Rb anti NG2 (Millipore # Ab5320), Rb anti MOG (Proteintech # 12690-1-ap), Rb anti Iba1 (Wako #016-20001) Gp anti VGlut1 (Millipore #AB5905, 1:2000), Gp anti VGlut2 (Millipore #AB2251 1:3000, 1:5000), Rb anti GluA1 (Millipore #AB1504, 1:400), Rb anti GluA2 (Millipore #AB1768-I 1:400), Ms anti Bassoon (Enzo #VAMP500, 1:500). All secondary antibodies were applied at 1:500 dilution.

The following mouse lines and antibody combinations were used:

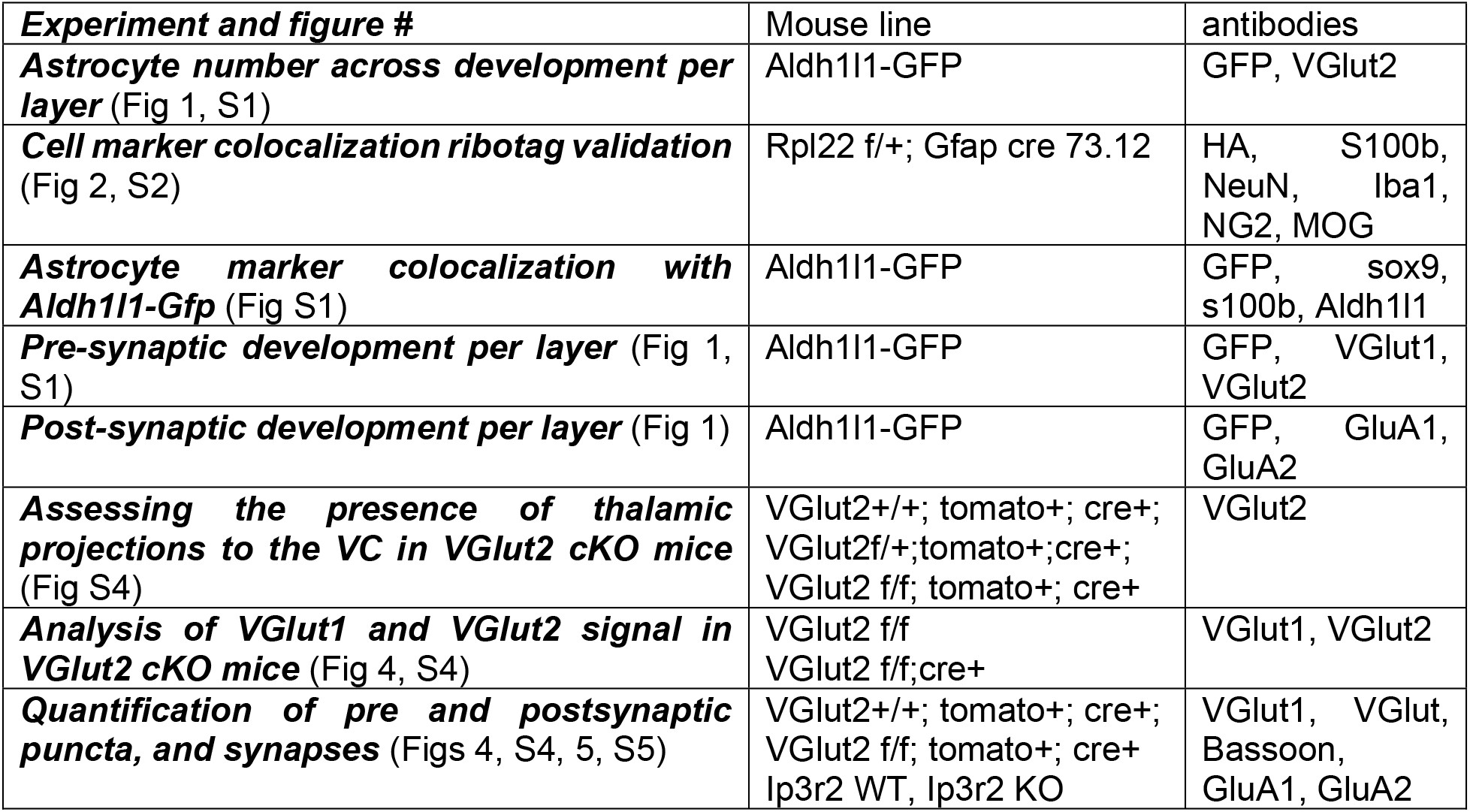

#### Single-molecule fluorescent in situ hybridization (smFISH)

The assay was performed using the RNAscope 2.5 HD—multiplex fluorescent Manual Assay kit (ACDbio #320850) using the manufacturer’s instructions modified for Fixed-frozen tissue. Briefly, slides containing brain sections were dried for 1 hr at −20C, then washed for 5 min in PBS at room temp, followed by brief wash (~1 min) in 100% Molecular Biology Grade Ethanol. The slides were then air-dried for 5 min, and incubated with appropriate pretreatment reagent at 40C. For P1-P7 – protease 3 30min; P14-P28 – protease 4 30 min. Slides were then briefly washed with PBS and incubated with target probes for 2 hrs at 40C, followed by 3 amplification steps and 1 detection step. Slides were mounted using the SlowFade Gold with DAPI mounting media (Life Tech #S36939) covered with 1.5 glass coverslip (Fisher #12544E) and sealed with clear nail polish.

All slides were either imaged within 1-2 days, or stored at −20C until imaging.

### IMAGING AND ANALYSIS

#### Fluorescent microscopy

All imaging was performed using an Axio Imager.Z2 fluorescent microscope (Zeiss) with the apotome module (apotome 2.0), and AxioCam HR3 camera (Zeiss) at 20x magnification. Tile images that contain the entire primary visual cortex (from pial surface to white matter tract) were acquired. Number of tiles adjusted to contain a similar area of the cortex at each developmental stage, typically 1-2(width) X 2-4 (depth) (pixel size 0.3X0.3 μm).

For developmental analysis of astrocyte numbers per layer (Fig 1, S1), presynaptic marker analysis during development (Fig 1, S1), VGlut2cKO validation (Fig 4, S4), and Aldh1l1-GFP mouse validation (Fig S1) - z stack images (3 slices, optical slice 1 μm) were obtained.

For Ribotag validation (Fig 2, S2) - Single plane images were obtained.

For In situ hybridization experiments (Fig 3, S3) - z stack images (7 slices, optical slice 1 μm) were obtained.

#### Confocal Microscopy

##### Developmental analysis of GluA1 and GluA2 expression

(Fig 1) slides were imaged using Zeiss LSM 700 confocal microscope at 63X magnification. An 1176X1176 pixel 2.7 um thick z stack image was obtained (pixel size 0.09X0.09X0.3 μm, 10 slices per 2.7 μm stack). In total 4 images were taken from each section to encompass all cortical layers. Layers 4-5 were combined into 1 image.

##### Imaging RORacre-tdTomato+ thalamic projections in the VC

(Fig S4) slides were imaged using Zeiss LSM 700 confocal microscope at 63X magnification. A 900 × 900 pixels 2.7 μm thick z stack image was obtained (xyz size 0.11X0.11X0.3 μm, 10 slices per 2.7 μm stack). Separate images were taken for Layer 1 and Layer 4.

##### Imaging synaptic proteins for synapse number analysis

(Figs 4, S4, 5, S5) slides were imaged using Zeiss LSM 880 confocal microscope at 63X magnification. For each section, 1420 × 920 pixels 3.5 μm thick z stack image was obtained (pixel size 0.08X0.08X0.39 μm; 10 slices per 3.5 μm stack). All images were from Layer 1. Example images show a single z plane from the same location in the stack for both genotypes.

In all cases, when comparing WT and KO per given experiment, slides were imaged on the same day using set exposure.

#### Image analysis

Image analysis was primarily done with ImageJ (NIH) or Imaris (Bitplane) software as described below for each section:

##### Astrocyte number across development per layer

(Fig 1, S1) was done on sections of Aldh1l1-GFP VC that were co-immunostained for VGlut2 using semi-automatic custom-made macro in ImageJ. For each image, maximal intensity projections were created, then each cortical layer was manually cropped based on DAPI and VGlut2 staining, and saved as a separate file. Then, a colocalization file was created using the “colocalization threshold” function to merge the colocalized cell marked by DAPI with the Aldh1l1-GFP signal to specifically select astrocytes. Colocalized objects were counted using the “analyze particles” function. Number of astrocytes in each layer was recorded for each developmental stage.

The high cell density at early ages made it impossible to use a similar method to count the total number of cells using the DAPI labels. Instead, 3 ROIs were created for each layer, and cells were counted manually within each ROI, using the “multi-point” tool in ImageJ. The total cell number in each layer was then extrapolated based on the total area measurement in each file.

##### Cell marker colocalization ribotag validation

(Fig 2, S2) was performed using FIJI (ImageJ). Thresholding was performed on the ribotag labeled image (stained with an anti-HA tag antibody) and the ‘Analyze Particles’ function was used with a minimum area of 20-40μm to automatically separate and quantify the total number of ribotag positive cells. The number of double-labeled ribotag and cell type antibody-positive cells were manually counted. This generated the proportion of ribotag positive cells that also label for the cell-specific marker.

##### Astrocyte marker colocalization with Aldh1l1-Gfp

(Fig S1) an ROI containing the entire depth of the cortex of a maximal intensity projection image, was cropped equally for each image. Labeled cells were counted manually using the “cell counter” plugin in ImageJ. Positively labeled cells were identified based on signal strength. For each file, 3 types of counts were made: the appropriate astrocyte marker positive cell number, Aldh1l1-GFP positive cell number and colocalized cell number.

##### Pre-synaptic development per layer

(Fig 1, S1) slides were immunostained for the presynaptic markers VGlut1 and VGlut2 as described above and the signal was analyzed with ImageJ. As above, for each file, maximal intensity projections were created, then layers were cropped out manually and saved as separate files. VGlut signal was thresholded in the same way for all images to contain all visible signal. The threshold area measurement was recorded for each file.

##### Post-synaptic development per layer

(Fig 1) slides were immunostained for the postsynaptic markers GluA1 and GluA2 as described above. Confocal z-stack images were analyzed using the Imaris software (Bitplane). GluA puncta number was calculated using the “spots” function, and mean intensity filter to select the positive puncta. All images were thresholded in the same way. To analyze specifically the GluA signal in the cell processes and not the soma, cell bodies labeled by DAPI were selected manually using the “create object” function. Then GluA puncta number that colocalized with cell bodies was found. Finally, cell body-related GluA1 puncta were subtracted from the total puncta number to obtain process-expressed GluAs.

##### Assessing the presence of thalamic projections to the VC in VGlut2 cKO mice

(Fig S4) confocal z-stack images were analyzed using Imaris (Bitplane). tdTomato positive processes were rendered using the “create object” function. All images were thresholded in the same way to select labeled processes. Total volume was calculated and compared between the experimental groups.

VGlut2 puncta were rendered using the “create spots” function, and mean intensity filter to threshold positive spots.

##### Analysis of VGlut1 and VGlut2 signal in VGlut2 cKO mice

(Fig 4, S4) images were analyzed using ImageJ as described for developmental presynaptic experiments.

##### Counting astrocytes in VGlut2 cKO and IP3R2 KO mice

(Figs S4, S5) smFISH in situ images (see below) were used to count astrocyte numbers within each layer. Counting was performed manually using the “cell counter” plugin in ImageJ. Astrocytes were identified by positive Glast probe signal.

##### Quantification of smFISH signal

(Figs 3, S3, 4, S4, 5, S5) was performed using a custom-made macro in ImageJ. Maximal intensity projection images of the visual cortex were manually cropped per layer and saved as individual files. Astrocytes were identified using the GFP signal in experiments with Aldh1l1-GFP mice (Fig3, S3); and Glast probe signal in all other experiments (Figs 4, S4, 5, S5). An ROI outline was then created around the cell body of the astrocyte, the probe of interest was then thresholded in the same way for all images, and the threshold area was recorded for each cell. We found that threshold area of the signal gave a more reliable and stable result than intensity measurements, which are not recommended by smFISH protocol (RNAscope by ACDbio). Due to the density of the signal in some cases, counting individual puncta was impossible.

##### Quantification of pre and postsynaptic puncta, and synapses

(Figs 4, S4, 5, S5) 3D z stack images were analyzed using Imaris software (Bitplane). Positive puncta of GluA1, GluA2, VGlut1, VGlut2, Bassoon, and tdTomato processed (FigS4) were selected by size and intensity by thresholding the images in the same way for each section. Then colocalization between each 2 pre-postsynaptic pairs was calculated. Puncta were considered colocalized if the distance between them was ≤ 0.5 μm (Blanco-Suarez et al., 2018; Farhy-Tselnicker et al., 2017). For experiments described in FigS4L-S, first colocalization between tdTomato and Bassoon was established, and cropped. The colocalized Bassoon-tdTomato puncta were then used to calculate colocalization with GluA1, or GluA2. Number of colocalized puncta was obtained and compared between the experimental groups. A minimum of 3 sections per mouse were imaged for each brain region, and the experiment was repeated in at least 5 WT and KO pairs. Example images show a single z plane from the same location in the stack for both genotypes.

### DATA PRESENTATION AND STATISTICAL ANALYSIS

All data is presented as either mean ± s.e.m, scatter with mean ± s.e.m, or scatter with range, as indicated in each figure legend. Statistical analysis was performed using Prism software (Graphpad). Multiple group comparisons were done using one-way Analysis of Variance (ANOVA) with post hoc Tukey’s or Dunn’s tests. Pairwise comparisons were done by T-test. When data did not pass the normal distribution test, multiple comparisons were done by Kruskal-Wallis ANOVA on ranks and pairwise comparisons were done with the Mann-Whitney Rank Sum test. P-value ≤ 0.05 was considered statistically significant. The sample sizes, statistical tests used and significance are presented in each figure and figure legend.

### DATA AVAILABILITY

The Ribotag data is available at GEO GSE161398 and glial snRNAseq at GSE163775.

