## Supplemental Figures for "Activity-Dependent Modulation of Synapse-Regulating Genes in Astrocytes"

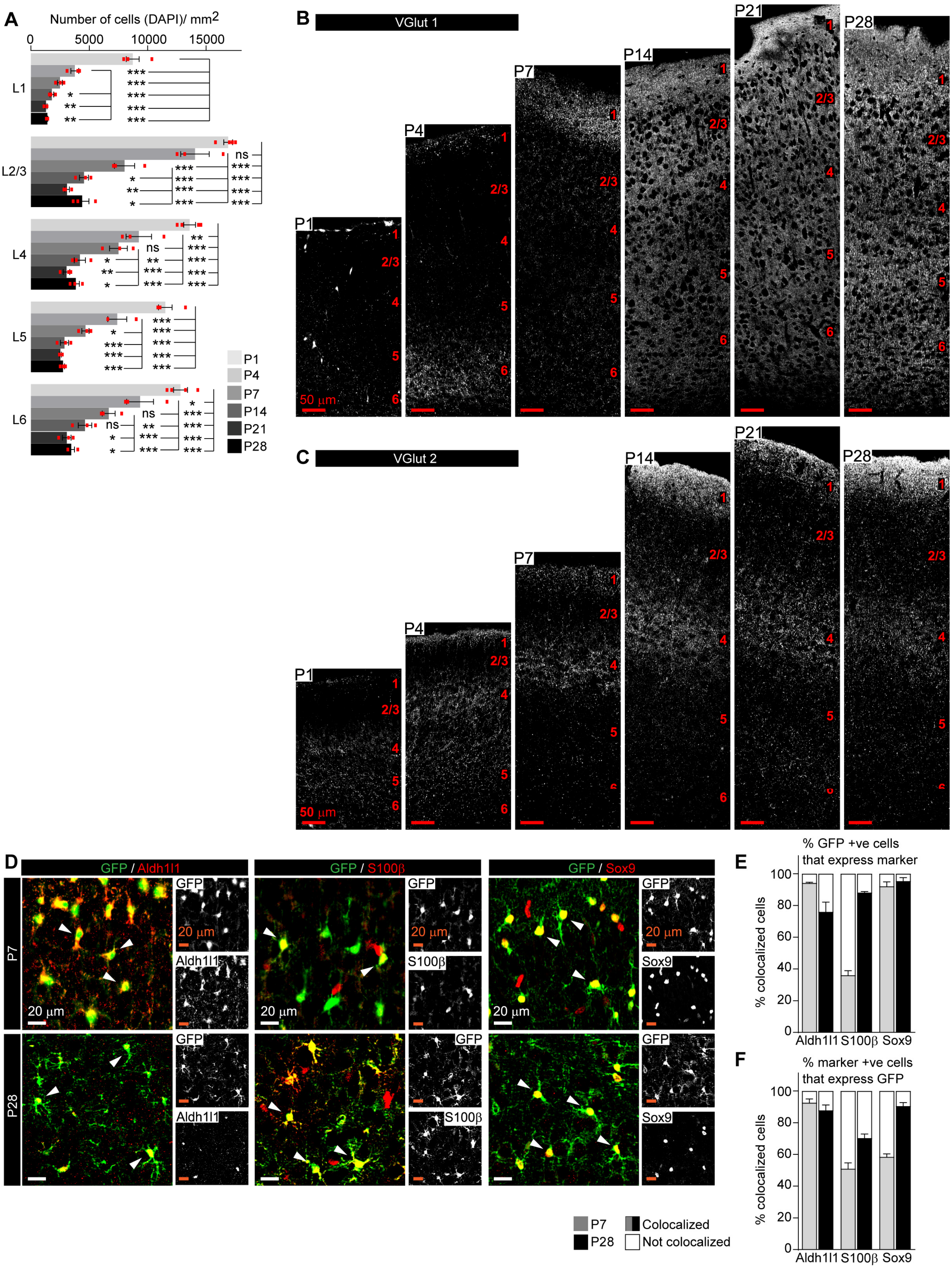
A
Number of cells (DAPI)/ mm2
0
5000
10000
15000
L1
L2/3
L4
L5
L6
ns
P1
P4
P7
P14
P21
P28
B
VGlut 1
50 μm
C
VGlut 2
50 μm
D
GFP / Aldh1l1
GFP / S100β
GFP / Sox9
P7
P28
GFP
Aldh1l1
S100β
Sox9
20 μm
E
% GFP +ve cells that express marker
% colocalized cells
Aldh1l1 S100β Sox9
F
% marker +ve cells that express GFP
% colocalized cells
Aldh1l1 S100β Sox9
P7
P28
Colocalized
Not colocalized

**Figure S1 (related to Figure 1). Development of astrocytes and synapses in the mouse visual cortex. A.** Quantification of Fig 1C, number of DAPI labeled cells per $mm^2$ of VC within each layer. N=3 mice/age. Graphs show mean ±s.e.m., red squares average of individual mouse. *P ≤ 0.05 **P<0.01, ***P<0.001, by one-way ANOVA comparing expression between time points within each layer. **B-C.** Example images of the entire span of the VC from Aldh1l1-GFP mice immunostained with antibody for Vglut1 (**B**), or VGlut2 (**C**) as indicated (white puncta), at time points analyzed. Scale bars = 50 µm. Layers indicated by numbers on the right of each panel in red. **D-F.** Aldh1l1-GFP signal colocalizes with astrocyte markers during development. **D.** Example images of the VC from Aldh1l1-GFP mice immunostained with antibodies for GFP (green), and astrocyte markers Aldh1l1, S100β, or Sox9 (as indicated, red) at the different time points as indicated. Inset is single channel image for each antibody. **E-F.** Quantification of D. **E.** Quantification of % of GFP +ve cells that are also immunopositive for the astrocyte marker demonstrating the majority of GFP +ve cells express astrocyte markers and are astrocytes. **F.** Quantification of % of astrocyte marker +ve cells that express GFP demonstrating that not all S100β and Sox9 +ve cells express GFP at P7. Data shows mean ± s.e.m. Arrowheads mark representative cells with colocalized GFP + marker. N=4 mice for P1; N=3 mice for P4-P28. Scale bars = 20 µm.

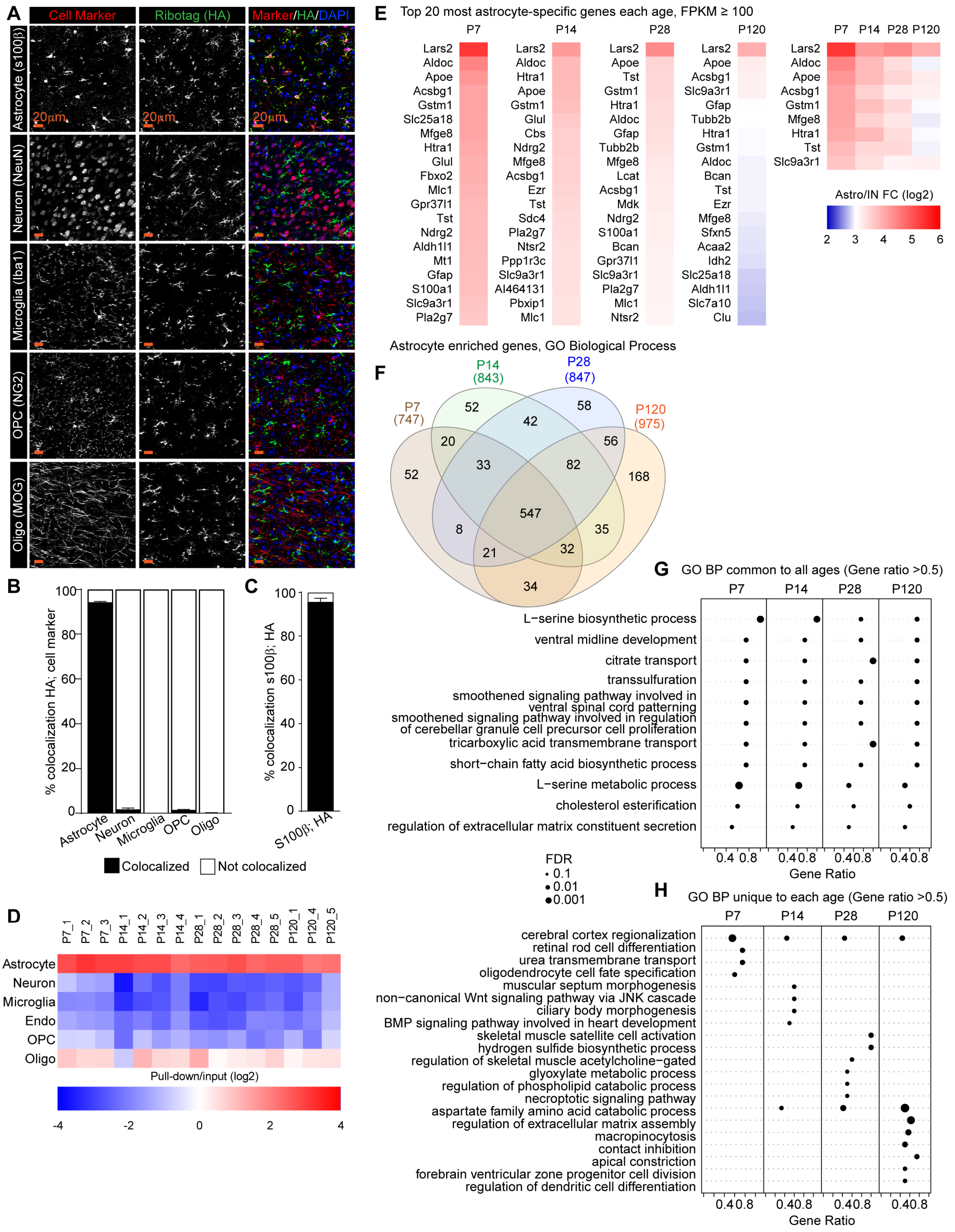
A
Cell Marker
Ribotag (HA)
Marker/HA/DAPI
Astrocyte (s100β)
Neuron (NeuN)
Microglia (Iba1)
OPC (NG2)
Oligo (MOG)
20µm
B
% colocalization HA; cell marker
Astrocyte
Neuron
Microglia
OPC
Oligo
C
% colocalization s100β; HA
S100β; HA
Colocalized
Not colocalized
D
P7_1 P7_2 P7_3 P14_1 P14_2 P14_3 P14_4 P28_1 P28_2 P28_3 P28_4 P28_5 P120_1 P120_4 P120_5
Astrocyte
Neuron
Microglia
Endo
OPC
Oligo
Pull-down/input (log2)
-4 -2 0 2 4
E
Top 20 most astrocyte-specific genes each age, FPKM ≥ 100
P7
Lars2 Aldoc Apoe Acsbg1 Gstm1 Slc25a18 Mfge8 Htra1 Glul Fbxo2 Mlc1 Gpr37l1 Tst Ndrg2 Aldh1l1 Mt1 Gfap S100a1 Slc9a3r1 Pla2g7
P14
Lars2 Aldoc Htra1 Apoe Gstm1 Glul Cbs Ndrg2 Mfge8 Acsbg1 Ezr Tst Sdc4 Pla2g7 Ntsr2 Ppp1r3c Slc9a3r1 AI464131 Pbxip1 Mlc1
P28
Lars2 Apoe Tst Gstm1 Htra1 Aldoc Gfap Tubb2b Mfge8 Lcat Acsbg1 Mdk Ndrg2 S100a1 Bcan Gpr37l1 Slc9a3r1 Pla2g7 Mlc1 Ntsr2
P120
Lars2 Apoe Acsbg1 Slc9a3r1 Gfap Tubb2b Htra1 Gstm1 Aldoc Bcan Tst Ezr Mfge8 Sfxn5 Acaa2 Idh2 Slc25a18 Aldh1l1 Slc7a10 Clu
P7 P14 P28 P120
Lars2 Aldoc Apoe Acsbg1 Gstm1 Mfge8 Htra1 Tst Slc9a3r1
Astro/IN FC (log2)
2 3 4 5 6
Astrocyte enriched genes, GO Biological Process
F
P7 (747)
P14 (843)
P28 (847)
P120 (975)
52 52 58 168 20 42 56 33 82 547 8 35 21 32 34
G
GO BP common to all ages (Gene ratio >0.5)
P7 P14 P28 P120
L−serine biosynthetic process
ventral midline development
citrate transport
transsulfuration
smoothened signaling pathway involved in ventral spinal cord patterning
smoothened signaling pathway involved in regulation of cerebellar granule cell precursor cell proliferation
tricarboxylic acid transmembrane transport
short−chain fatty acid biosynthetic process
L−serine metabolic process
cholesterol esterification
regulation of extracellular matrix constituent secretion
0.4 0.8
Gene Ratio
FDR
0.1
0.01
0.001
H
GO BP unique to each age (Gene ratio >0.5)
P7 P14 P28 P120
cerebral cortex regionalization
retinal rod cell differentiation
urea transmembrane transport
oligodendrocyte cell fate specification
muscular septum morphogenesis
non−canonical Wnt signaling pathway via JNK cascade
ciliary body morphogenesis
BMP signaling pathway involved in heart development
skeletal muscle satellite cell activation
hydrogen sulfide biosynthetic process
regulation of skeletal muscle acetylcholine−gated
glyoxylate metabolic process
regulation of phospholipid catabolic process
necroptotic signaling pathway
aspartate family amino acid catabolic process
regulation of extracellular matrix assembly
macropinocytosis
contact inhibition
apical constriction
forebrain ventricular zone progenitor cell division
regulation of dendritic cell differentiation
0.4 0.8
Gene Ratio

**Figure S2 (related to Figure 2). Determination of the astrocyte transcriptome across visual cortex development. A-C.** Immunostaining for the HA-tag and cell specific markers to determine cell-type expression of tagged ribosomes in P28 VCs from Rpl22-HA$^{f/+;}$ Gfap-cre 73.12 mice. Left panels, cell marker; middle panels, HA; right panels, merge with DAPI to mark nuclei. **B-C.** Quantification of A. **B.** Colocalization of HA with each cell-specific marker, expressed as % of marker +ve cells, demonstrates majority of HA +ve cells are astrocytes. **C.** Quantification of number of astrocytes (s100β +ve) that are also HA positive, demonstrates majority of astrocytes express HA-tagged ribosomes. N=4 mice s100β, NeuN, Iba1; 3 mice Mog; 2 mice Ng2. Bar graphs mean ± s.e.m. Scale bars = 20 μm. **D.** Analysis of cell-specific genes in HA-pulldown/input samples (log2) from RNA sequencing, demonstrates enrichment for astrocyte genes and depletion of other cells in all samples (N=3 at P7, 4 at P14, 5 at P28, 3 at P120. For statistical comparisons 3xP120 samples published in (Boisvert et al., 2018) were added to increase the power of the analysis). **E.** Heatmaps of top 20 most astrocyte-specific genes at each timepoint, as well as those in the top 20 at all timepoints, represented as FC of astrocyte/input (log2). **F-H.** GO terms analysis with String db of Biological process (BP) in astrocyte enriched genes at each time point. **F.** Venn diagram showing overlap in GO terms at each age. **G.** Dot plot of GO terms common to all time points, gene ratio >0.5. **H.** Dot plot of GO terms unique to each age, with gene ratio >0.5. Size of dot is FDR, position on the axis is gene ratio. See also Table S2B.

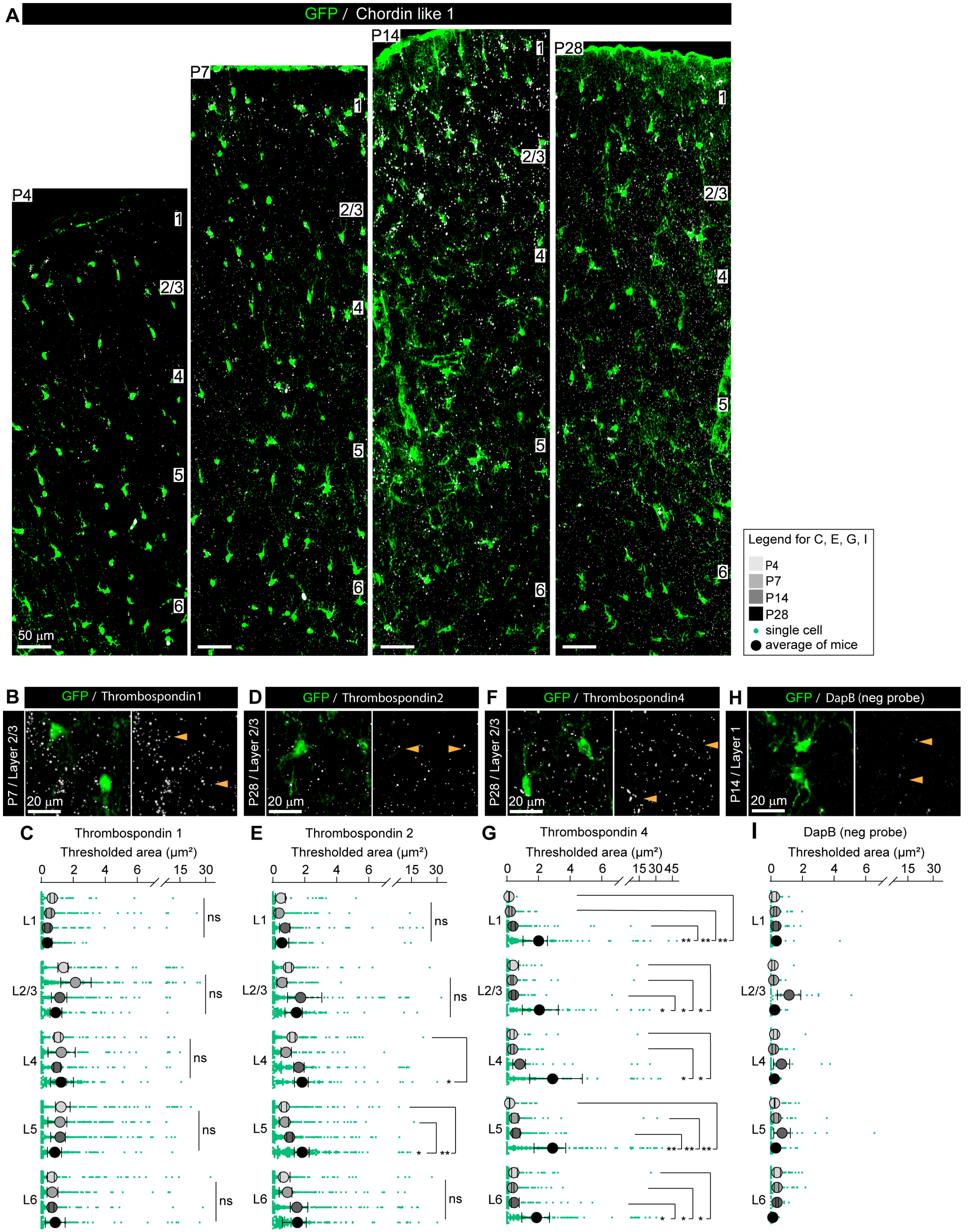

A
GFP / Chordin like 1
P4
P7
P14
P28
1
2/3
4
5
6
50 µm
Legend for C, E, G, I
P4
P7
P14
P28
single cell
average of mice
B
GFP / Thrombospondin1
P7 / Layer 2/3
20 µm
D
GFP / Thrombospondin2
P28 / Layer 2/3
20 µm
F
GFP / Thrombospondin4
P28 / Layer 2/3
20 µm
H
GFP / DapB (neg probe)
P14 / Layer 1
20 µm
C
Thrombospondin 1
Thresholded area (µm²)
0 2 4 6 15 30
L1
L2/3
L4
L5
L6
ns
E
Thrombospondin 2
Thresholded area (µm²)
0 2 4 6 15 30
L1
L2/3
L4
L5
L6
ns
*
**
G
Thrombospondin 4
Thresholded area (µm²)
0 2 4 6 15 30 45
L1
L2/3
L4
L5
L6
*
**
I
DapB (neg probe)
Thresholded area (µm²)
0 2 4 6 15 30
L1
L2/3
L4
L5
L6

**Figure S3 (related to Figure 3). Synapse regulating genes in astrocytes show differential spatio-temporal expression. A.** Example images of the entire span of the VC showing Chrdl1 mRNA (white) in astrocytes (green) at each age and layer as labeled. **B, D, F, H.** Example images showing Thbs1, Thbs2, Thbs4 and DapB mRNA (white) in astrocytes (green) at each age and layer as labeled. Merged panel on the left, single channel probe panel on the right. **C, E, G, I.** Quantification of B, D, F, H respectively. **C.** No difference in expression of Thbs1 at any age or layer. **E.** Thbs2 expression is increased at P28 in L4-5. **G.** Thbs4 expression is increased at P28 in all layers. **I.** DapB negative probe expression level showing detection limit for probe mRNA expression analysis. Arrowheads in single channel panel mark astrocyte cells on the left. Data presented as scatter with mean + range. Green dots are mRNA signal measured in individual astrocyte. Large circles colored according to time points are average signal. N=3 mice/age, n=~50-350 astrocytes/ per age total; n=~20-80 astrocytes for DapB neg probe (H-I) (average and statistical analysis is calculated based on N=3 i. e. data per mouse). Scale bars = 20 µm. *P ≤ 0.05 **P<0.01, ns: not significant by one-way ANOVA comparing expression between time points within each layer; see also Tables S3A-B.

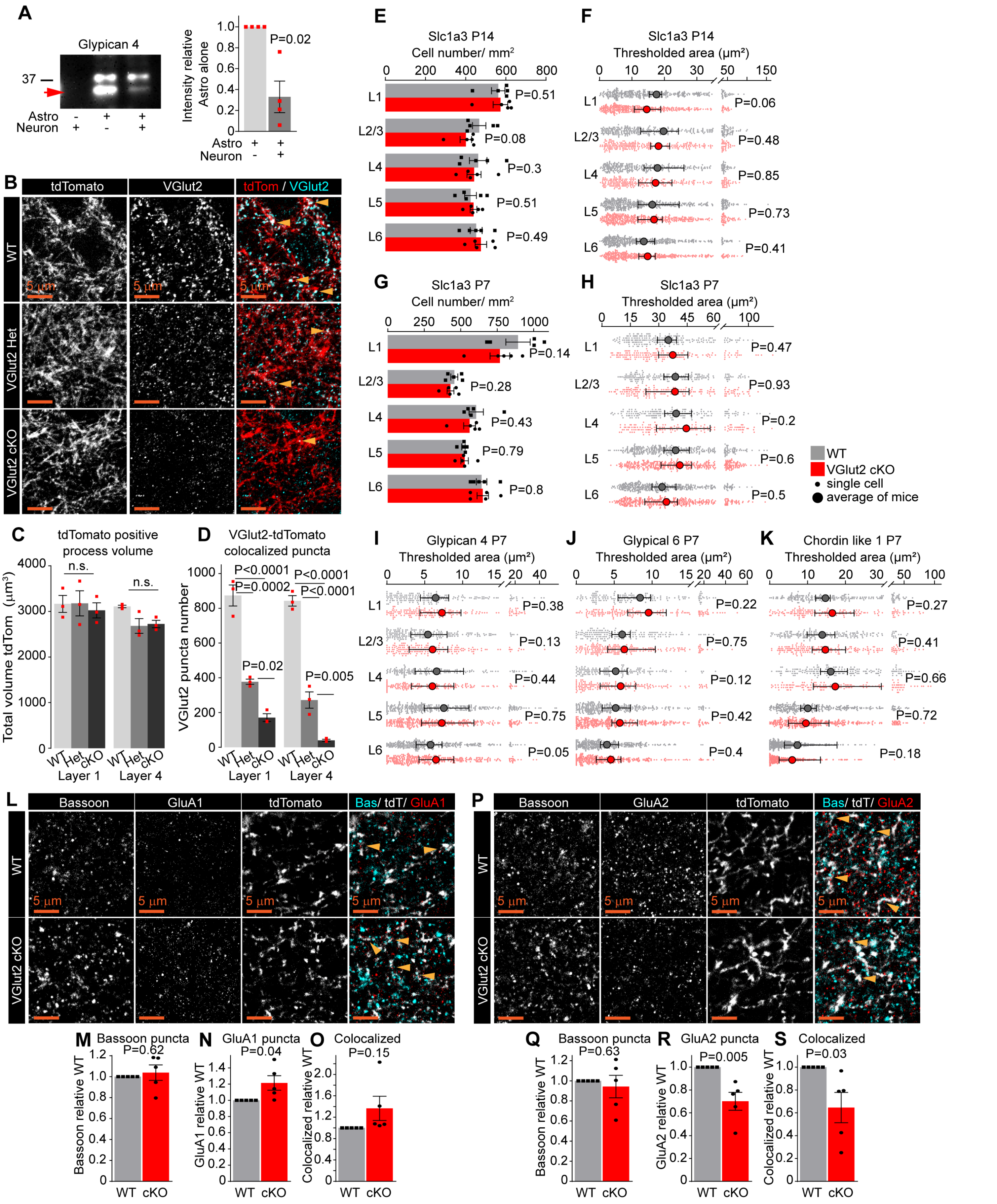
A
Glypican 4
37
Astro - + +
Neuron + - +
Intensity relative Astro alone
P=0.02
B
tdTomato
VGlut2
tdTom / VGlut2
WT
VGlut2 Het
VGlut2 cKO
5 µm
C
tdTomato positive process volume
Total volume tdTom (µm³)
n.s.
WT Het cKO
Layer 1
Layer 4
D
VGlut2-tdTomato colocalized puncta
VGlut2 puncta number
P<0.0001
P=0.0002
P<0.0001
P<0.0001
P=0.02
P=0.005
E
Slc1a3 P14
Cell number/ mm²
P=0.51
P=0.08
P=0.3
P=0.51
P=0.49
F
Slc1a3 P14
Thresholded area (µm²)
P=0.06
P=0.48
P=0.85
P=0.73
P=0.41
G
Slc1a3 P7
Cell number/ mm²
P=0.14
P=0.28
P=0.43
P=0.79
P=0.8
H
Slc1a3 P7
Thresholded area (µm²)
P=0.47
P=0.93
P=0.2
P=0.6
P=0.5
WT
VGlut2 cKO
single cell
average of mice
I
Glypican 4 P7
Thresholded area (µm²)
P=0.38
P=0.13
P=0.44
P=0.75
P=0.05
J
Glypical 6 P7
Thresholded area (µm²)
P=0.22
P=0.75
P=0.12
P=0.42
P=0.4
K
Chordin like 1 P7
Thresholded area (µm²)
P=0.27
P=0.41
P=0.66
P=0.72
P=0.18
L1
L2/3
L4
L5
L6
L
Bassoon
GluA1
tdTomato
Bas/ tdT/ GluA1
P
Bassoon
GluA2
tdTomato
Bas/ tdT/ GluA2
M
Bassoon puncta
P=0.62
Bassoon relative WT
N
GluA1 puncta
P=0.04
GluA1 relative WT
O
Colocalized
P=0.15
Colocalized relative WT
Q
Bassoon puncta
P=0.63
Bassoon relative WT
R
GluA2 puncta
P=0.005
GluA2 relative WT
S
Colocalized
P=0.03
Colocalized relative WT
WT cKO

**Figure S4 (related to Figure 4). Neuronal activity tunes astrocyte expression of synapse-regulating genes. A.** Western blot showing Gpc4 secretion from astrocytes is decreased in the presence of neurons. Example blot on the left, quantification on the right. Data presented as mean ± s.e.m, normalized to control condition, astrocytes cultured alone. N=4 independent cultures. P by t-test, value on the plot. Red arrow indicates Gpc4 signal at ~36 KDa. **B-D.** LGN axons innervate the VC in the absence of VGlut2. **B.** Example images of L4 VC showing thalamic axons (tdTomato, red) and VGlut2 (cyan) presence in WT, VGlut2 cHet, and VGlut2 cKO mice as labeled. **C-D.** Quantification of B. **C.** No difference in volume of tdTomato +ve LGN axons between the genotypes in either L1 or L4 as indicated. **D.** VGlut2 puncta associated with tdTomato +ve processes are reduced in VGlut2 cHet, and VGlut2 cKO compared to WT. Plots show mean signal ± s.e.m. Squares are average of signal in each mouse. N=3 mice/genotype. Scale bar = 5 μm. P by One-Way ANOVA comparing genotypes within each layer, value on the plot. n.s: not significant. **E, G.** Number of astrocytes (Slc1a3 positive cells) is unaltered in VGlut2 cKO mice at P14 or P7. Data presented as mean ± s.e.m. squares and circles each mouse. N=5 mice/ genotype. P by paired t-test, value on the plot. **F, H.** Glast (Slc1a3) mRNA expression levels are unaltered in VGlut2 cKO mice at P14 or P7. Data presented as scatter with mean + range, large circles are average signal from data per mouse. Grey or red dots are mRNA signal measured in individual astrocyte in WT and VGlut2 cKO respectively. N=5 mice/genotype, n=~150-450 astrocytes/ per age total (average and statistical analysis is calculated based on N=5 i.e. per mouse). **I-K.** mRNA expression of Gpc4, Gpc6 and Chrdl1 is unaltered in VGlut2 cKO mice at P7. Data presented as scatter with mean + range, large circles average signal from data per mouse. Grey or red dots are mRNA signal measured in individual astrocyte in WT and VGlut2 cKO respectively. N=5 mice/genotype, n=~150-450 astrocytes/ per age total (average and statistical analysis is calculated based on N=5 i.e. per mouse). Statistical analysis by paired t-test, P value on each plot. See also Tables S4A-B. **L-O.** Increase in GluA1 protein levels in L1 of the VC in VGlut2 cKO mice. **L.** Example images from WT (top) and cKO (bottom), Bassoon in cyan, GluA1 in red, and tdTomato positive LGN axon in white. **M, N, O.** Quantification of Bassoon (number of puncta within the tdTomato positive processes), GluA1 and colocalized puncta respectively, normalized to WT. **P-S.** Decrease in GluA2 protein levels and colocalization between GluA2 and Bassoon in L1 of the VC in VGlut2 cKO mice. **P.** Example images from WT (top) and cKO (bottom), Bassoon in cyan, GluA2 in red, and tdTomato positive thalamic axon in white. **Q, R, S.** Quantification of Bassoon (number of puncta within the tdTomato positive processes), GluA2 and colocalized puncta respectively, normalized to WT. In L-S data presented as mean ± s.e.m, squares and circles are mean fold change of each mouse. N=5 mice/ genotype. Arrowheads mark representative colocalized puncta. Scale bar = 5 μm. Statistical analysis by t-test, P value on each plot.

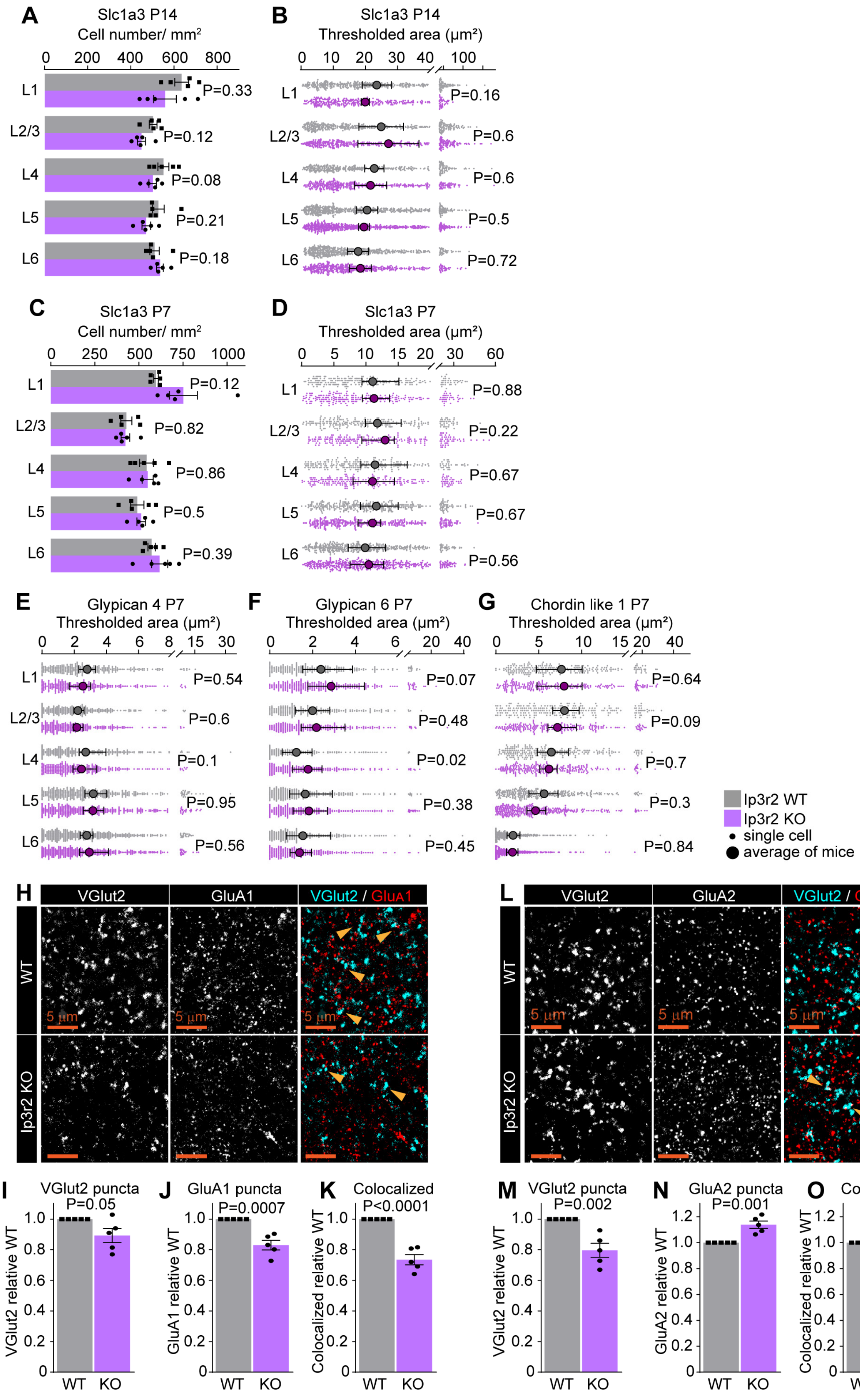

A
Slc1a3 P14
Cell number/ mm²
B
Slc1a3 P14
Thresholded area (µm²)
C
Slc1a3 P7
Cell number/ mm²
D
Slc1a3 P7
Thresholded area (µm²)
E
Glypican 4 P7
Thresholded area (µm²)
F
Glypican 6 P7
Thresholded area (µm²)
G
Chordin like 1 P7
Thresholded area (µm²)
Ip3r2 WT
Ip3r2 KO
single cell
average of mice
H
VGlut2
GluA1
VGlut2 / GluA1
WT
Ip3r2 KO
5 µm
L
VGlut2
GluA2
VGlut2 / GluA2
I
VGlut2 puncta
P=0.05
J
GluA1 puncta
P=0.0007
K
Colocalized
P<0.0001
M
VGlut2 puncta
P=0.002
N
GluA2 puncta
P=0.001
O
Colocalized
P=0.21

**Figure S5 (related to Figure 5). Astrocyte calcium signaling regulates expression of synapse-regulating genes. A, C.** Number of astrocytes (Slc1a3 positive cells) is unaltered in Ip3r2 KO mice at P14 or P7. Data presented as mean ± s.e.m. squares and circles are each mouse. N=5 mice/ genotype. P by paired t-test, value on the plot. **B, D.** Glast (Slc1a3) mRNA expression levels are unaltered in Ip3r2 KO mice at P14 or P7. Data presented as scatter with mean + range. Grey or purple dots are mRNA signal measured in individual astrocyte in WT, and KO respectively. Large circles are average signal. N=5 mice/genotype, n=~50-350 astrocytes/ per age total (see also Table S4; average and statistical analysis is calculated based on N=5 i.e. per mouse). **E-G.** mRNA expression of Gpc4 (**E**), and Chrdl1 (**G**) is unaltered in Ip3r2 KO mice at P7. Gpc6 (**F**) expression is increased in L4. Data presented as scatter with mean + range. Grey or purple dots are mRNA signal measured in individual astrocyte in WT, and KO respectively. Large circles are average signal. N=5 mice/genotype, n=~150-450 astrocytes / per age total (average and statistical analysis is calculated based on N=5 i.e. per mouse). Statistical analysis by paired t-test, P value on each plot. See also Tables 4A-B. **H-K.** Decrease in VGlut2, GluA1 protein levels, and colocalization between GluA1 and VGlut2 in L1 of the VC in Ip3r2 KO mice. **H.** Example images from WT (top) and KO (bottom), VGlut2 in cyan and GluA1 in red. **I, J, K.** Quantification of VGlut2, GluA1 and colocalized puncta respectively, normalized to WT. **L-O.** Decrease in VGlut2, and increase GluA2 protein levels, with no change in colocalization between GluA2 and VGlut2 in L1 of the VC in Ip3r2 KO mice. **L.** Example images from WT (top) and KO (bottom), VGlut2 in cyan and GluA2 in red. **M, N, O.** Quantification of VGlut2, GluA2 and colocalized puncta respectively, normalized to WT. In H-O data presented as mean ± s.e.m, squares and circles each mouse. N=5 mice/genotype. Arrowheads mark representative colocalized puncta. Scale bar = 5 μm. Statistical analysis by t-test, P value on each plot.

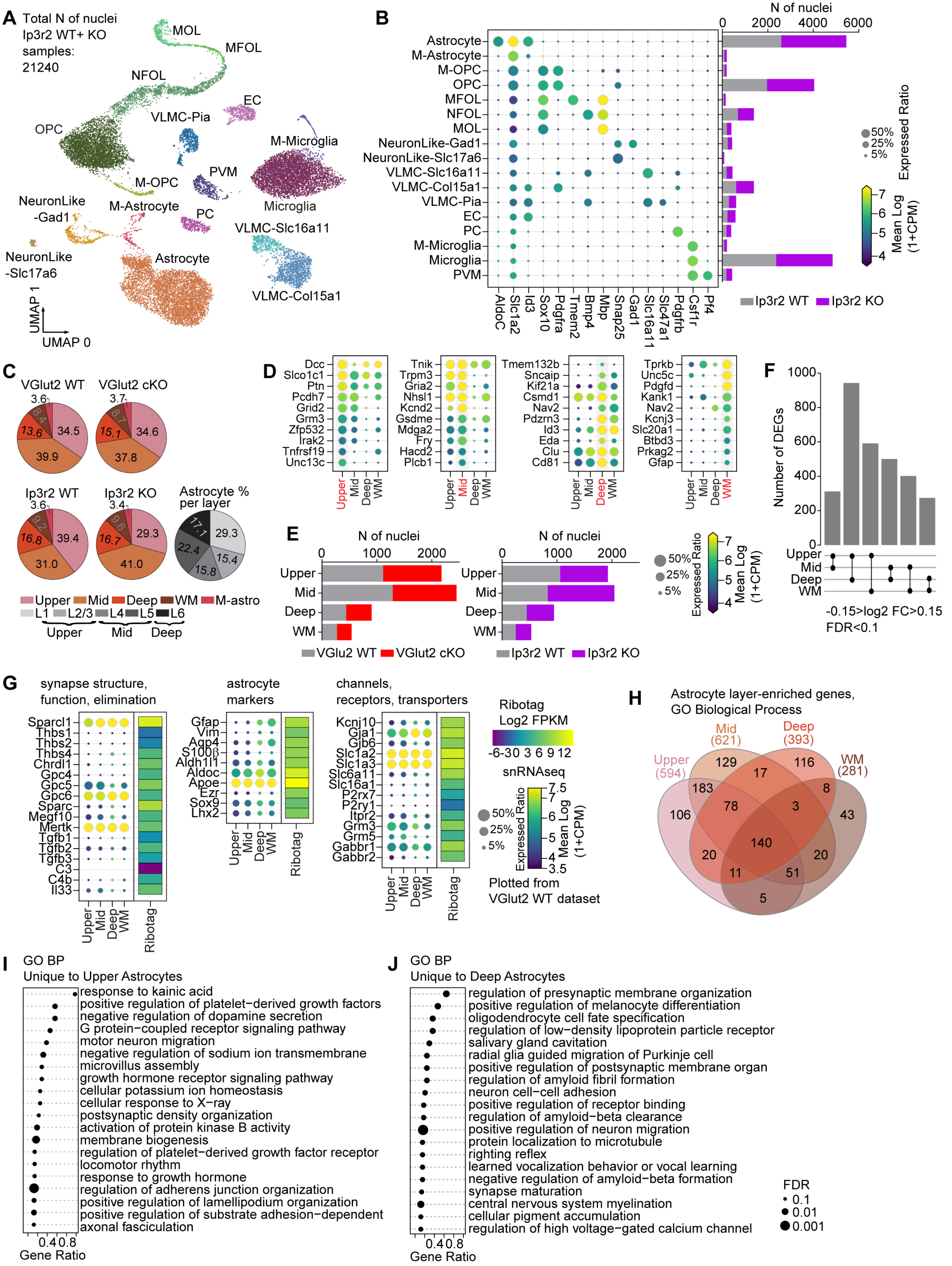
A
Total N of nuclei
Ip3r2 WT+ KO
samples:
21240
MOL
MFOL
NFOL
EC
VLMC-Pia
OPC
M-Microglia
M-OPC
PVM
Microglia
NeuronLike
-Gad1
M-Astrocyte
PC
VLMC-Slc16a11
NeuronLike
-Slc17a6
Astrocyte
VLMC-Col15a1
UMAP 1
UMAP 0
B
N of nuclei
0
2000
4000
6000
Astrocyte
M-Astrocyte
M-OPC
OPC
MFOL
NFOL
MOL
NeuronLike-Gad1
NeuronLike-Slc17a6
VLMC-Slc16a11
VLMC-Col15a1
VLMC-Pia
EC
PC
M-Microglia
Microglia
PVM
AldoC
Slc1a2
Id3
Sox10
Pdgfra
Tmem2
Bmp4
Mbp
Snap25
Gad1
Slc16a11
Slc47a1
Pdgfrb
Csf1r
Pf4
Ip3r2 WT
Ip3r2 KO
50%
25%
5%
Expressed Ratio
7
6
5
4
Mean Log
(1+CPM)
C
VGlut2 WT
VGlut2 cKO
3.6
8.4
13.6
34.5
39.9
3.7
8.7
15.1
34.6
37.8
Ip3r2 WT
Ip3r2 KO
3.6
9.2
16.8
39.4
31.0
3.4
9.6
16.7
29.3
41.0
Astrocyte %
per layer
17.1
29.3
22.4
15.4
15.8
Upper
Mid
Deep
WM
M-astro
L1
L2/3
L4
L5
L6
Upper
Mid
Deep
D
Dcc
Slco1c1
Ptn
Pcdh7
Grid2
Grm3
Zfp532
Irak2
Tnfrsf19
Unc13c
Tnik
Trpm3
Gria2
Nhsl1
Kcnd2
Gsdme
Mdga2
Fry
Hacd2
Plcb1
Tmem132b
Sncaip
Kif21a
Csmd1
Nav2
Pdzrn3
Id3
Eda
Clu
Cd81
Tprkb
Unc5c
Pdgfd
Kank1
Nav2
Kcnj3
Slc20a1
Btbd3
Prkag2
Gfap
Upper
Mid
Deep
WM
E
N of nuclei
0
1000
2000
Upper
Mid
Deep
WM
VGlu2 WT
VGlut2 cKO
N of nuclei
0
1000
2000
Upper
Mid
Deep
WM
Ip3r2 WT
Ip3r2 KO
50%
25%
5%
Expressed Ratio
7
6
5
4
Mean Log
(1+CPM)
F
1000
800
600
400
200
0
Number of DEGs
Upper
Mid
Deep
WM
-0.15>log2 FC>0.15
FDR<0.1
G
synapse structure,
function, elimination
Sparcl1
Thbs1
Thbs2
Thbs4
Chrdl1
Gpc4
Gpc5
Gpc6
Sparc
Megf10
Mertk
Tgfb1
Tgfb2
Tgfb3
C3
C4b
Il33
Upper
Mid
Deep
WM
Ribotag
astrocyte
markers
Gfap
Vim
Aqp4
S100β
Aldh1l1
Aldoc
Apoe
Ezr
Sox9
Lhx2
Upper
Mid
Deep
WM
Ribotag
channels,
receptors, transporters
Kcnj10
Gja1
Gjb6
Slc1a2
Slc1a3
Slc6a11
Slc16a1
P2rx7
P2ry1
Itpr2
Grm3
Grm5
Gabbr1
Gabbr2
Upper
Mid
Deep
WM
Ribotag
Ribotag
Log2 FPKM
-6-30 3 6 9 12
snRNAseq
50%
25%
5%
Expressed Ratio
7.5
7
6
5
4
3.5
Mean Log
(1+CPM)
Plotted from
VGlut2 WT dataset
H
Astrocyte layer-enriched genes,
GO Biological Process
Mid
(621)
Deep
(393)
Upper
(594)
WM
(281)
129
17
116
183
8
78
3
106
43
140
20
20
11
51
5
I
GO BP
Unique to Upper Astrocytes
response to kainic acid
positive regulation of platelet−derived growth factors
negative regulation of dopamine secretion
G protein−coupled receptor signaling pathway
motor neuron migration
negative regulation of sodium ion transmembrane
microvillus assembly
growth hormone receptor signaling pathway
cellular potassium ion homeostasis
cellular response to X−ray
postsynaptic density organization
activation of protein kinase B activity
membrane biogenesis
regulation of platelet−derived growth factor receptor
locomotor rhythm
response to growth hormone
regulation of adherens junction organization
positive regulation of lamellipodium organization
positive regulation of substrate adhesion−dependent
axonal fasciculation
0.4
0.8
Gene Ratio
J
GO BP
Unique to Deep Astrocytes
regulation of presynaptic membrane organization
positive regulation of melanocyte differentiation
oligodendrocyte cell fate specification
regulation of low−density lipoprotein particle receptor
salivary gland cavitation
radial glia guided migration of Purkinje cell
positive regulation of postsynaptic membrane organ
regulation of amyloid fibril formation
neuron cell−cell adhesion
positive regulation of receptor binding
regulation of amyloid−beta clearance
positive regulation of neuron migration
protein localization to microtubule
righting reflex
learned vocalization behavior or vocal learning
negative regulation of amyloid−beta formation
synapse maturation
central nervous system myelination
cellular pigment accumulation
regulation of high voltage−gated calcium channel
0.4
0.8
Gene Ratio
FDR
0.1
0.01
0.001

**Figure S6 (related to Figure 6). Unbiased determination of astrocyte layer-enriched genes. A.** UMAP clustering of different cell types identified in the NeuN negative population of the combined samples from Ip3r2 WT and KO mice. 17 clusters were identified including the 3 main types of glia: astrocytes, oligodendrocytes and microglia, as well as endothelial cells, and two sub-types of neurons (Abbreviations are: M-astrocyte – mitotic astrocyte; M-OPC – mitotic oligodendrocyte precursor cell; OPC – oligodendrocyte precursor cell; MFOL - myelin forming oligodendrocyte; NFOL - newly formed oligodendrocyte; MOL - mature oligodendrocyte; VLMC- vascular and leptomeningeal cell; EC – Endothelial cell; PC - pericyte; PVM - perivascular macrophage). **B.** Expression level of select marker genes for each cluster. Circle size denotes expression ratio (percent cells expressing the gene), color represents expression level (in Log2 CPM). Bar chart on the right is number of cells identified for each genotype as labeled. Similar cell numbers were identified for Ip3r2 WT and KO groups for each cluster. **C.** Pie charts as labeled showing percent astrocytes for each identified cluster out of total astrocytes. Monochrome pie chart is percent of astrocytes out of total astrocytes observed within each cortical layer in histological experiments (Fig1). **D.** Unbiased clustering analysis identified 4 sub-populations of astrocytes in the P14 VC, using Ip3r2 WT dataset. Populations annotated to Upper, Mid, Deep and White matter types following comparison with published datasets. Panels show select list of 10 genes that are highly expressed in each population as indicated. Size of the circle is expression ratio; color is expression level (log2 CPM). **E.** Similar cell numbers were identified for Vglut2 WT and cKO; and for Ip3r2 WT and KO groups for each astrocyte group. **F.** Pairwise comparison identified ~300-900 differentially expressed genes between populations in Ip3r2 WT mice. Criteria for DEG selection: Log2 FC between -0.15 and 0.15; FDR <0.1. See also Table S5B. **G.** Comparison of expression levels of astrocyte markers, function and synapse-related genes identified in bulk RNAseq (Fig2) with the snRNAseq dataset shows overall positive correlation between expression levels obtained by both methods. For snRNAseq panels, size of the circle is expression ratio; color is expression level (log2 CPM). For bulk RNAseq heatmaps, log2 FPKM is shown. **H-J.** GO terms analysis with String db of Biological process of layer group-specific genes. **H.** Venn diagram showing overlap in GO terms between layer groups. **I-J.** Dot plots of GO terms unique to Upper (**I**), and Deep (**J**) astrocytes, showing divergent GO BP enrichment. Size of dot is FDR, position on the axis is gene ratio. See also Table S5C.

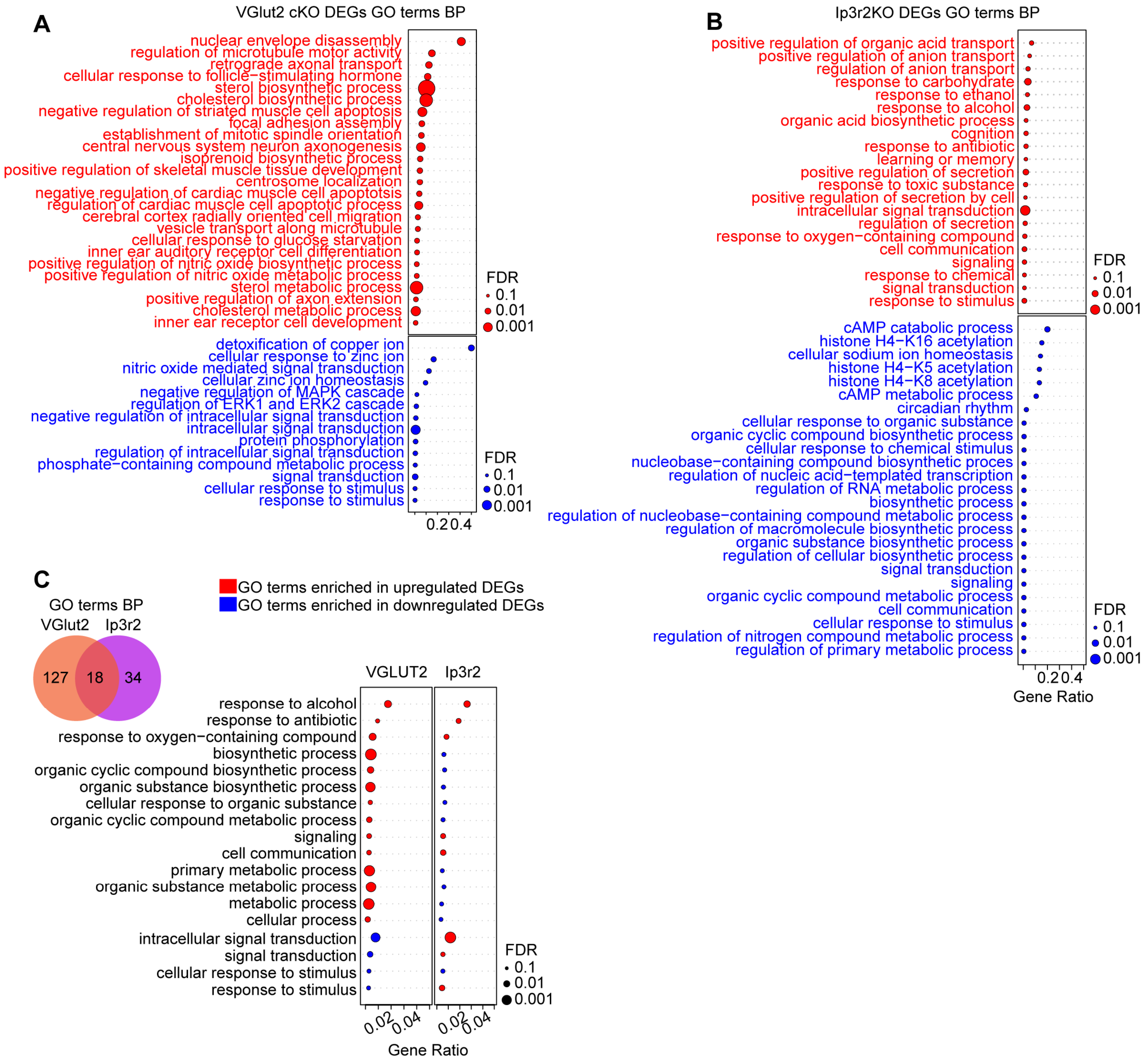
A
VGlut2 cKO DEGs GO terms BP
nuclear envelope disassembly
regulation of microtubule motor activity
retrograde axonal transport
cellular response to follicle−stimulating hormone
sterol biosynthetic process
cholesterol biosynthetic process
negative regulation of striated muscle cell apoptosis
focal adhesion assembly
establishment of mitotic spindle orientation
central nervous system neuron axonogenesis
isoprenoid biosynthetic process
positive regulation of skeletal muscle tissue development
centrosome localization
negative regulation of cardiac muscle cell apoptotsis
regulation of cardiac muscle cell apoptotic process
cerebral cortex radially oriented cell migration
vesicle transport along microtubule
cellular response to glucose starvation
inner ear auditory receptor cell differentiation
positive regulation of nitric oxide biosynthetic process
positive regulation of nitric oxide metabolic process
sterol metabolic process
positive regulation of axon extension
cholesterol metabolic process
inner ear receptor cell development
FDR
0.1
0.01
0.001
detoxification of copper ion
cellular response to zinc ion
nitric oxide mediated signal transduction
cellular zinc ion homeostasis
negative regulation of MAPK cascade
regulation of ERK1 and ERK2 cascade
negative regulation of intracellular signal transduction
intracellular signal transduction
protein phosphorylation
regulation of intracellular signal transduction
phosphate−containing compound metabolic process
signal transduction
cellular response to stimulus
response to stimulus
FDR
0.1
0.01
0.001
0.2
0.4
B
Ip3r2KO DEGs GO terms BP
positive regulation of organic acid transport
positive regulation of anion transport
regulation of anion transport
response to carbohydrate
response to ethanol
response to alcohol
organic acid biosynthetic process
cognition
response to antibiotic
learning or memory
positive regulation of secretion
response to toxic substance
positive regulation of secretion by cell
intracellular signal transduction
regulation of secretion
response to oxygen−containing compound
cell communication
signaling
response to chemical
signal transduction
response to stimulus
FDR
0.1
0.01
0.001
cAMP catabolic process
histone H4−K16 acetylation
cellular sodium ion homeostasis
histone H4−K5 acetylation
histone H4−K8 acetylation
cAMP metabolic process
circadian rhythm
cellular response to organic substance
organic cyclic compound biosynthetic process
cellular response to chemical stimulus
nucleobase−containing compound biosynthetic proces
regulation of nucleic acid−templated transcription
regulation of RNA metabolic process
biosynthetic process
regulation of nucleobase−containing compound metabolic process
regulation of macromolecule biosynthetic process
organic substance biosynthetic process
regulation of cellular biosynthetic process
signal transduction
signaling
organic cyclic compound metabolic process
cell communication
cellular response to stimulus
regulation of nitrogen compound metabolic process
regulation of primary metabolic process
FDR
0.1
0.01
0.001
0.2
0.4
Gene Ratio
C
GO terms BP
VGlut2
Ip3r2
127
18
34
GO terms enriched in upregulated DEGs
GO terms enriched in downregulated DEGs
VGLUT2
Ip3r2
response to alcohol
response to antibiotic
response to oxygen−containing compound
biosynthetic process
organic cyclic compound biosynthetic process
organic substance biosynthetic process
cellular response to organic substance
organic cyclic compound metabolic process
signaling
cell communication
primary metabolic process
organic substance metabolic process
metabolic process
cellular process
intracellular signal transduction
signal transduction
cellular response to stimulus
response to stimulus
FDR
0.1
0.01
0.001
0.02
0.04
0.02
0.04
Gene Ratio

**Figure S7 (related to Figure 7). Global astrocyte gene expression changes following silencing of neuronal or astrocyte activity. A-C.** GO terms analysis with String db of Biological process in DEGs from the VGlut2 cKO model (**A**), Ip3r2 KO model (**B**), and common DEGs to both models (**C**). Blue indicates terms enriched in downregulated DEGs, red terms enriched in upregulated DEGs. Size of dot is FDR, position on the axis is gene ratio. See also Table S6D.
